# A Meta-Analysis of the Converging Effects of Different Classes of Antipsychotics on the Frontal Cortex Transcriptome in Laboratory Rodents and Non-Human Primates

**DOI:** 10.64898/2026.06.24.734301

**Authors:** Mubashshir Ra’eed Bhuiyan, Megan Hastings Hagenauer, Eva Geoghegan, Elizabeth Flandreau, Stanley Watson, Huda Akil

## Abstract

**Background:** Psychotic illnesses are among the most debilitating classes of psychiatric disorders, requiring targeted and effective treatment strategies. Although antipsychotics are the primary pharmacological therapy for psychosis, their full range of effects remain unclear, including effects within the frontal cortex, a brain region linked structurally and functionally to psychotic disorders.

**Methods:** To examine the effects of antipsychotic treatment on the frontal cortex, we conducted a meta-analysis of publicly available rodent (rat, mice) transcriptional profiling datasets (microarray, RNA-Seq). Five datasets (GSE45229, GSE93918, GSE2547, GSE4031.1, GSE66275) were identified within the Gemma database using pre-specified search terms and inclusion/exclusion criteria (date: 7/7/2024), yielding differential expression results for eight drug vs. control comparisons (collective *n*=68). A random-effects meta-analysis model was fit to the log2 fold changes for each gene, and p-values adjusted for false discovery rate (FDR), with follow-up analyses exploring robustness, heterogeneity, and publication bias. To increase the power and generalizability of our findings, an exploratory meta-analysis was also run incorporating antipsychotic effects from both rodents and nonhuman primates (collective *n*=101), and compared to findings from individuals with schizophrenia.

**Results:** Our meta-analysis yielded stable estimates for 12,190 genes, identifying 63 genes that were differentially expressed following antipsychotic treatment (“DEGs”, FDR<0.05). Differential expression included genes important for serotonergic and cholinergic signalling, and was enriched within pathways linked to oligodendrocyte development and myelination, physiological and cellular stress responses, and cardiovascular function. An exploratory meta-analysis combining rodent and nonhuman primate results confirmed these observations and yielded additional findings (117 DEGs total). Comparisons with human post-mortem findings suggested that some schizophrenia-related gene expression may instead reflect antipsychotic treatment.

**Conclusion:** Further validation is necessary, but our findings suggest that antipsychotics may assist in the regulation of specific structural and functional changes within the frontal cortex linked to psychotic disorders.

**Graphical Abstract:** 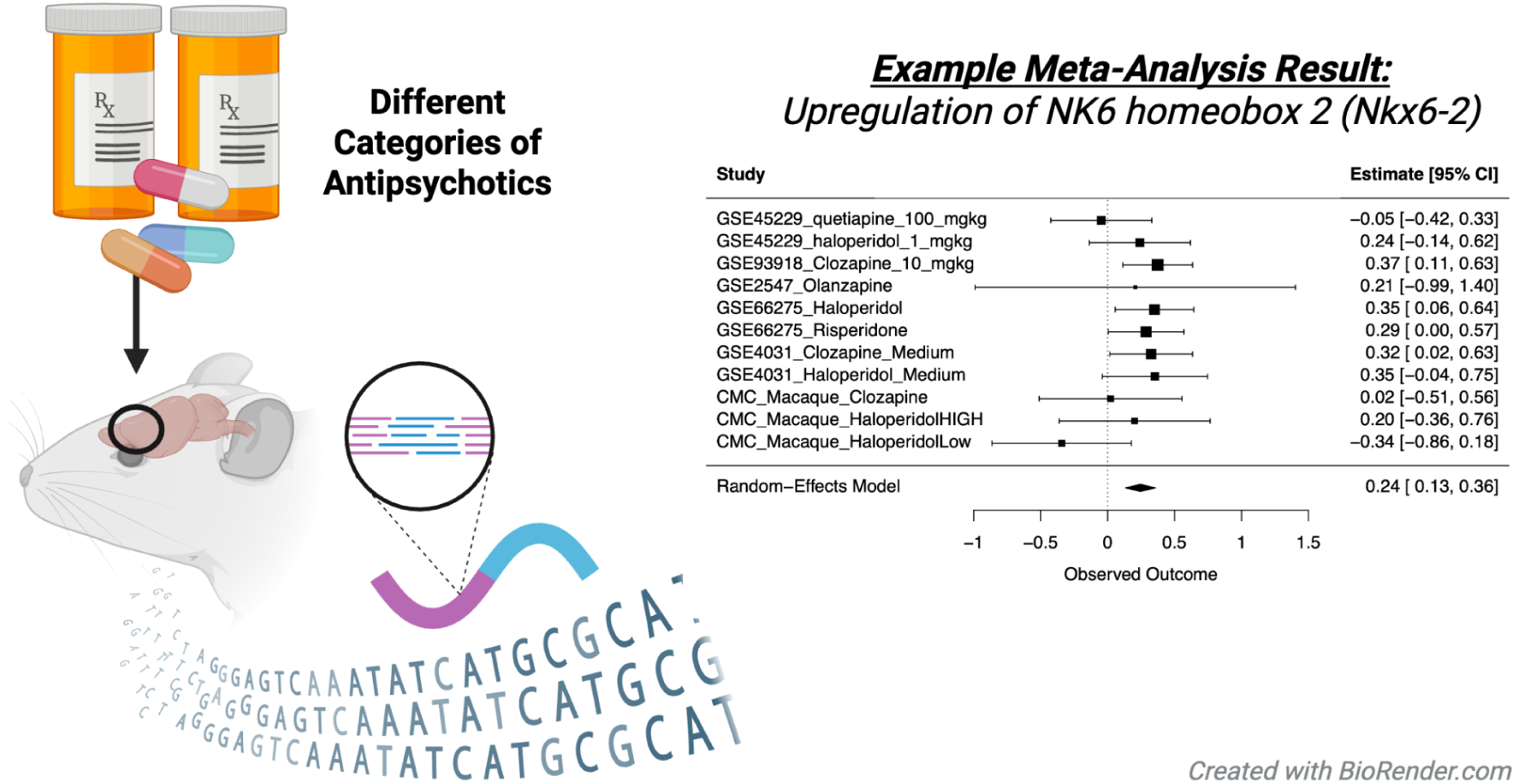

**Key Points:**

- Psychosis is treated using two broad types of antipsychotic medication: first-generation (typical) and second-generation (atypical).
- Understanding the congruent effects of different types of antipsychotics on regions important for psychosis, such as the frontal cortex, can highlight essential mechanisms.
- A meta-analysis of public transcriptional profiling datasets identified genes and functional gene sets that are differentially expressed across antipsychotic categories.

## Introduction

Psychosis is a broad term used to describe a mental state characterized by an inability to properly distinguish tangible reality from thought (Lieberman & First, 2018), as occurs in mental disorders such as schizophrenia, bipolar disorder, schizoaffective disorder, and postpartum psychosis (Arciniegas, 2015; Lieberman & First, 2018). Schizophrenia, the most common psychotic disorder, affects approximately 1 out of 300 people worldwide (∼24 million people) (*GBD Results*, n.d.; Huhn et al., 2019), and is both expensive and straining, with a high median societal cost per patient (Christensen et al., 2020). Given the heavy burden and clinical significance of psychotic disorders, targeted and effective treatment strategies are essential to adequately manage symptoms and improve patient outcomes.

The standard treatment for psychosis is antipsychotic medication (Huhn et al., 2019). Antipsychotic drugs are commonly separated into two broad categories: first-generation (typical) and second-generation (atypical). Typical antipsychotics function primarily by antagonizing dopamine D_2_ receptors. They are effective at reducing positive symptoms of psychosis (*e.g.,* hallucinations, disorganized behavior, thought disorders), but frequently associated with extrapyramidal side effects (EPS), such as tremors, rigidity, and tardive dyskinesia (Julayanont & Suryadevara, 2021; Siafis et al., 2023). In contrast, atypical antipsychotics modulate both dopaminergic and serotonergic neurotransmission, including antagonizing serotonin 5-HT_2_A receptors (H. Y. Meltzer, 2013). They can be effective against both positive and negative symptoms (Grinchii & Dremencov, 2020) and carry a lowered risk of EPS (H. Meltzer & Massey, 2011; Romeo et al., 2023), making them the preferred first line treatment (Briles et al., 2012; Fabrazzo et al., 2022). Despite these differences, typical and atypical antipsychotics may not be fully distinguishable drug classes, due to overlapping receptor targets and inconsistent subjective efficacy (Leucht et al., 2024).

Although standard treatment, the full range of antipsychotic effects on the brain remains unclear, requiring further research to evaluate their full mechanisms of action (Bowling & Santini, 2016; Girgis et al., 2008). These mechanisms may involve the frontal cortex and its subregions, such as the prefrontal cortex (PFC), which is a critical node within a network of brain regions dysregulated in psychotic disorders (Chopra et al., 2025). The PFC is responsible for core executive functions, including working memory, cognitive flexibility, inhibitory control, emotional regulation, and decision-making (D. T. Jones & Graff-Radford, 2021). Functional and structural abnormalities within the PFC are linked to psychosis, such as altered myelination, response to oxidative stress, and neurotransmission (Bilecki & Maćkowiak, 2023). Converging evidence from structural and functional imaging, as well as post-mortem studies, suggests that the PFC serves as a key center within a broad neural circuit that includes the hippocampus and thalamus that is critical to schizophrenia pathogenesis (Morris, 2026). Moreover, antipsychotic treatment non-response has been associated with PFC structural and functional deficits, including disconnectivity with other brain regions (Bilecki & Maćkowiak, 2023; D. T. Jones & Graff-Radford, 2021), in a manner that may contribute to symptom persistence. Therefore, clarifying the effects of antipsychotics on the PFC may help identify functional pathways alleviating psychotic symptoms.

To characterize the effects of antipsychotics on the frontal cortex, we conducted a meta-analysis of publicly available transcriptional profiling datasets. Transcriptional profiling technologies, such as RNA sequencing (RNA-seq) and microarray, quantify gene expression across the genome, providing broad biological insight. However, drawing meaningful conclusions from individual datasets can be difficult due to small sample sizes and technical artifacts. Meta-analysis can reduce these limitations, drawing on a larger set of data to identify consistent patterns with greater statistical power (Olivas-Bernal et al., 2026). To identify pathways affected across drug categories, we included datasets using a wide variety of antipsychotic treatments, improving the generalizability of our results. We focused on studies using laboratory animals (mice, rats), as medication effects in human post-mortem data are often confounded by illness presentation and severity. To ensure adequate statistical power for our initial analysis, we included datasets from all regions of the frontal cortex, acknowledging that future research will be necessary to explore regional specificity and species differences (Preuss & Wise, 2022). To increase the power and generalizability of our findings, we then ran an exploratory meta-analysis incorporating antipsychotic effects from both rodents and nonhuman primates, and compared our results to findings from individuals with schizophrenia. We hope that examining differential expression induced by a wide variety of antipsychotics across the greater frontal cortex region will help identify novel directions for future research and provide a useful reference database to improve the interpretation of human post-mortem data from psychotic subjects.

## Methods

### General

Detailed methods for the meta-analysis and associated R code (R v.4.4.1; RStudio v2024.12.1.563 (Posit Team, 2025; R Core Team, 2024)) can be found at: https://github.com/raeedbhuiyan/Antipsychotic_Meta-Analysis. This meta-analysis was completed as part of the *Brain Data Alchemy Project*, a collective effort to improve the reliability and generalizability of transcriptional profiling findings using meta-analyses of public datasets. This guided effort uses a standardized pipeline for dataset identification, inclusion/exclusion, and meta-analysis (protocol for 2024: (M. Hagenauer et al., 2024), example validation: (Geoghegan et al., 2025; Rhoads et al., 2025; Xiong et al., 2026), *not preregistered*), leveraging the large scale data curation, preprocessing, and analysis efforts of the *Gemma* database (Lim et al., 2021; Zhou & Stephens, 2012). *Gemma* uses a standardized pipeline that includes dataset alignment to an updated genome, removal of outlier samples, removal of genes with minimal variance in expression, and manual curation for common issues such as batch effects. Differential expression is calculated using the *limma* or *limma-voom* pipeline, with statistical output available for the full model (omnibus) and individual contrasts (Ritchie et al., 2015; Smyth, 2005). To date, *Gemma* holds a collection of over 19,000 re-analyzed transcriptional profiling datasets (Lim et al., 2021), sourced from public repositories like the NCBI Gene Expression Omnibus (GEO) and Sequence Read Archive (SRA) (Barrett et al., 2013).

### Dataset identification and initial filtering

A strict search strategy was used to identify potential datasets, following the *Brain Data Alchemy Project* protocol (M. Hagenauer et al., 2024). The focus of the meta-analysis was on the effect of pharmaceutical treatments that target symptoms of psychosis (*i.e.,* antipsychotic medications). Gemma was briefly browsed to confirm whether relevant datasets were available to determine project viability. Relevant search terms were then generated using a comprehensive compilation of all publicly available antipsychotic medications as per the United State Food and Drug Administration (FDA) and *Rxnorm* database (Christian et al., 2012; *RxClass*, n.d.). Brand names as well as generic names were included in the final search term list, as well as synonyms and permutations of the term “antipsychotic”. These terms were then converted to formal search syntax (**Figure 1**).

**Figure 1.**
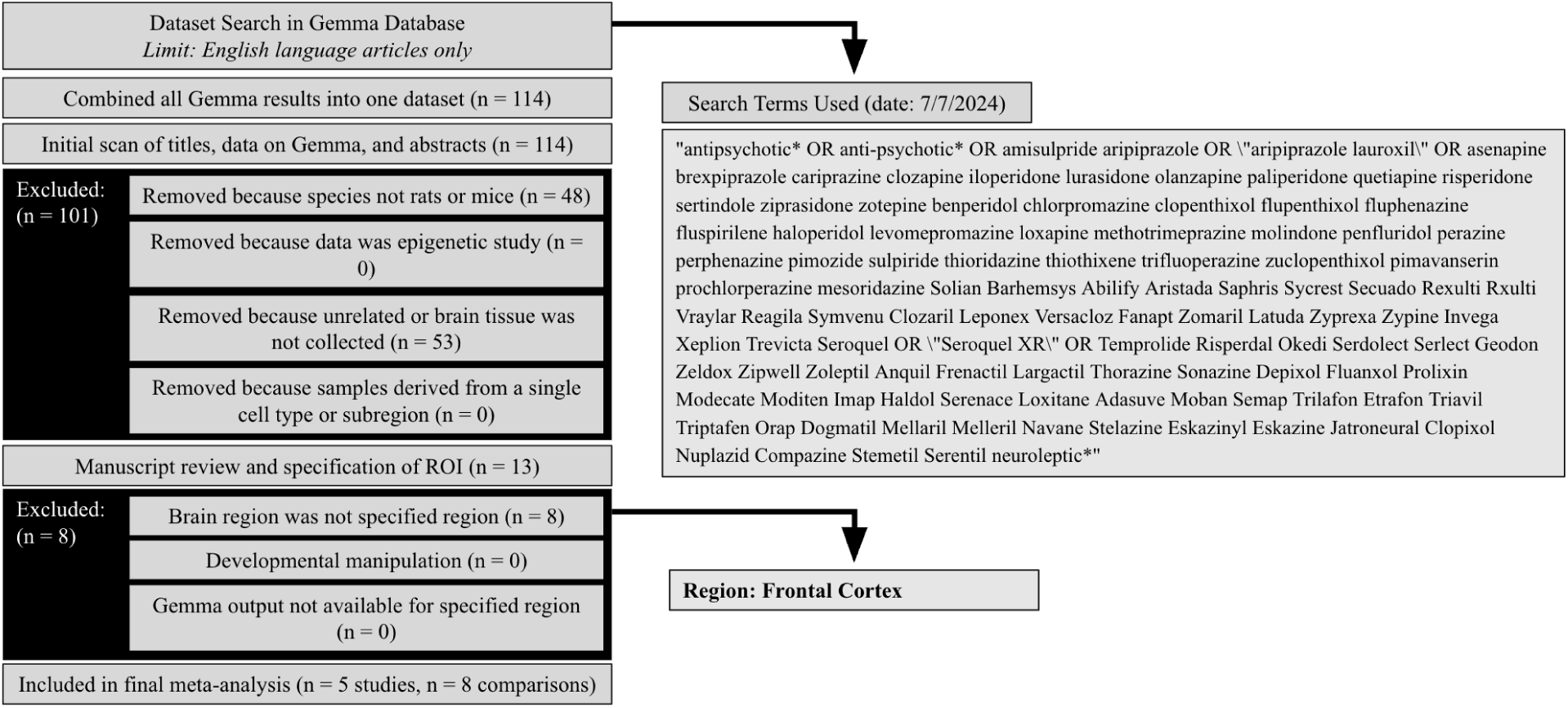
PRISMA Diagram of Dataset Search and Inclusion. We utilized pre-specified search terms to narrow down potentially viable transcriptional profiling datasets of antipsychotic effects. Initial Gemma results were compiled into one dataset in which the titles, abstracts, and metadata for the datasets were scanned and filtered for the pre-specified inclusion/exclusion criteria. Datasets were excluded that were not derived from rodent models or brain tissue. Remaining datasets’ manuscripts were reviewed in the secondary screening step, including a detailed review of the metadata on Gemma and associated publications, followed by a finalized selection of the brain tissue region of interest (ROI) based on the availability of datasets (ROI = frontal cortex).

The scope of the search encompassed publicly available transcriptional profiling datasets (gene expression microarray or RNA-Sequencing) in the *Gemma* database. The initial search yielded 114 potential datasets (search date: 7/7/2024). Following this search procedure, the NIH GEO Database was also later manually browsed to confirm that there were no additional applicable datasets that had not yet been curated by Gemma (search date: 2/26/2026).

The metadata for the potential datasets, including titles, abstracts, and dataset characteristics, were screened using pre-specified inclusion/exclusion criteria. During this initial screen, 48 datasets were excluded because the species were not the rodent models selected for inclusion (rats: *Rattus norvegicus*, mice: *Mus musculus*). Another 53 datasets were excluded because brain tissue was not collected. The remaining datasets met our initial inclusion criteria: 1) adult subjects, 2) RNA extracted from bulk-dissected brain tissue instead of a particular cell type or small subregion, 3) profiling the full transcriptome or a large representation of the full transcriptome and not a subpopulation of transcripts (*e.g.,* Chip-Seq, miRNA, TRAP-Seq, etc.), 4) no critical metadata issues, including duplication/overlap with another record or critical missing information. Reasons for dataset exclusion were documented by the first author (MRB) and reviewed and approved by a second researcher (MHH).

### Secondary filtering and selection of brain region of interest (ROI)

The remaining 13 datasets were filtered following a detailed review of the metadata and published methods. We selected the frontal cortex as the brain region of interest (ROI) for the meta-analysis due to the relative availability of datasets; 5 of 13 datasets were specific to, or included, the frontal cortex. The remaining eight datasets were derived from other brain regions, including the striatum, Ammon’s horn, and whole brain. Final dataset selection was conducted by the first author (MRB) and reviewed and approved by co-authors (EG, MHH). The complete dataset characteristics for the five datasets that fulfilled all inclusion/exclusion criteria (GSE45229, GSE93918, GSE2547, GSE66275, and GSE4031.1) are provided in **Table 1**, along with the sample sizes for the subset of the dataset representing the ROI.

**Table 1.** Overview of Included Datasets. “Dataset ID” refers to the Gene Expression Omnibus (GEO) accession number for each rodent dataset included in the planned meta-analysis and the research consortium responsible for each of the non-human primate and primate datasets used in the exploratory comparisons. The “sample size n (ROI)” is the total sample size for the brain region of interest. “The sample size n (subset)” is the subsetted sample size for the treatment vs. control comparisons included in our analysis. The “Species” column specifies the species and strain (when known) for the subjects in the dataset. The “Sex” column refers to the sex of the subjects. The “Author” column lists the first author of the publication associated with the study. The “Year” column reflects the publication date for the publication associated with the study. The “Platform” column indicates the transcriptional profiling technology used. The “Tissue” column notes the brain region sampled. The “Treatment Duration” column notes the duration of antipsychotic administration prior to tissue collection. The “Treatment (dose): n” column specifies the treatment groups in the study, including the antipsychotic medication and dosage, when known, and their respective sample sizes. All experimental details outline characteristics of the samples included within the transcriptional profiling study. [Note: Dataset number 5 (GSE4031.1) does not have an associated publication. The given author is the contributor listed within the dataset’s GEO submission. The given year is the year the dataset was made publicly available on GEO.]

| Dataset ID | n (ROI) | n (subset) | Species | Sex | Author | Year | Platform | Tissue | Treatment Duration | Treatment (dose): n |
| --- | --- | --- | --- | --- | --- | --- | --- | --- | --- | --- |
| **Planned Rodent Meta-Analysis:** |  |  |  |  |  |  |  |  |  |  |
| GSE45229 | 10 | 6 | Mus musculus (CD-1 mice) | Male | Kondo et al. | 2013 | Microarray (Affymetrix) | Frontal cortex | 28 days | - Quetiapine (100mg/kg): n=2<br>- Haloperidol (1mg/kg): n=2<br>- Vehicle: n=2 |
| GSE93918 | 12 | 12 | Mus musculus (wild type and HDAC2 knockout) | Male | Ibi et al. | 2017 | RNA-seq (Illumina) | PFC | 21 days | - Clozapine (10mg/kg): n=6<br>- Vehicle: n=6 |
| GSE2547 | 8 | 8 | Rattus norvegicus (Sprague–Dawley albino rats) | Male | Fatemi et al. | 2006 | Microarray (Affymetrix) | Frontal cortex | 21 days | - Olanzapine (2mg/kg): n=4<br>- Vehicle: n=4 |
| GSE66275 | 15 | 15 | Rattus norvegicus (Sprague–Dawley albino rats) | Male | Lanz et al. | 2019 | Microarray (Affymetrix) | Frontal cortex | 21 days | - Haloperidol (0.25 mg/kg): n=5<br>- Risperidone (5mg/kg): n=5<br>- Vehicle: n=5 |
| GSE4031.1 | 63 | 27 | Rattus norvegicus | Unk | Colantuoni | 2006 | Microarray (Affymetrix) | Frontal cortex | “Chronic” | - Clozapine (medium dose): n=9<br>- Haloperidol (medium dose): n=9<br>- Reference: n=9 |
| **Exploratory Comparisons:** |  |  |  |  |  |  |  |  |  |  |
| Common Mind Consortium | 33 | 33 | Macaca mulatta | Both | Schulmann et al. | 2023 | RNA-Seq | DLPFC | 6 months | - Clozapine (5.2 mg/kg): n=9<br>- Haloperidol (0.14 mg/kg): n=10<br>- Haloperidol (4 mg/kg): n=7<br>- Placebo: n=7 |
| Eli Lilly and Company | 42 | 30 | Macaca fascicularis | Unk | Martin et al. | 2015 | Microarray (Affymetrix) | PFC | Antipsychotic treatment: 28 days<br>Psychosis model (PCP): 42 days | - Haloperidol (1.5 mg/kg): n=6<br>- Olanzapine (0.75 mg/kg): n=6<br>- Vehicle: n=6<br>- PCP (1-2 mg/kg): n=6<br>- Olanzapine (0.75 mg/kg) + PCP (1-2 mg/kg): n=6 |
| Common Mind Consortium | 604 | 201 (Subset with Toxicology) | Homo sapiens | Both | Schulman et al. | 2023 | RNA-Seq (Illumina) | DLPFC | Unknown Except Toxicology at TOD | - Antipsychotic+ at TOD: n=65<br>- Antipsychotic- at TOD: n=23<br>- Control: n=113 |
| Common Mind Consortium | 604 | 557 | Homo sapiens | Both | Gandal et al. | 2018 | RNA-Seq (Illumina) | DLPFC | Treatment Status Unknown | - Schizophrenia: n=262<br>- Control: n=295 |
| Psych ENCODE | 700 | 452 | Homo sapiens | Both | Gandal et al. | 2018 | Microarray (Meta-Analysis) | Cortex | Treatment Status Unknown | - Schizophrenia: n=159<br>- Control: n=293 |

### Selection of treatment comparisons of interest

From these datasets, we selected relevant antipsychotic vs. control comparisons (statistical contrasts). For GSE93918, the comparison of interest was treatment for 21 days of clozapine vs. vehicle (Ibi et al., 2017). For GSE2547, the comparison of interest was treatment for 21 days of olanzapine vs. vehicle (Fatemi et al., 2006). For GSE66275, the comparisons of interest were treatment for 21 days of haloperidol vs. vehicle or treatment for 21 days of risperidone vs. vehicle (Lanz et al., 2019). For GSE45229 the comparison of interests were treatment for 28 days of quetiapine vs. vehicle or treatment for 28 days of haloperidol vs. vehicle (Kondo et al., 2013). However, the full sample size included two dosages of each treatment — 1 mg kg^−1^ and 0.3 mg kg^−1^ haloperidol and 100 mg kg^−1^ and 10 mg kg^−1^ quetiapine. Within the original publication, only the high dosage groups (1 mg kg^−1^ haloperidol and 100 mg kg^−1^ quetiapine) were considered representative of effective clinical dosages, with lower dosage conditions included “for reference purposes” (Kondo et al., 2013), so we excluded them from the meta-analysis. For GSE4031.1, no associated publication was available, and the dataset metadata provided limited contextual background, simply stating that “chronic” administration of clozapine was delivered at dose “low”, “medium”, and “high”, and “chronic” administration of haloperidol delivered at dose “low”, “medium”, and “high”. The “reference substance role” did not specify whether vehicle treatment was used as the control condition. To reduce heterogeneity within the meta-analysis, we opted to utilize only the medium-dose chronic haloperidol vs. reference comparison and medium-dose chronic clozapine vs. reference comparison. **Table 1** provides the sample sizes for the final comparisons of interest used in the meta-analysis (“n (subset)”).

### Individual dataset quality control and reprocessing

The preprocessed gene expression data from each of the selected datasets was accessed in Gemma to confirm that data fell into a typical range for log2 RNA-Seq gene expression (between −5 to 12) or log2 microarray gene expression (4 to 15). GSE4031.1 had a highly restricted range, suggesting miscalculation, and required manual reprocessing. GSE66275 also required reprocessing, as the record in Gemma did not include differential expression output specific to the frontal cortex, instead including “Organism part” (striatum, Ammon’s horn, and frontal cortex) as a variable in the differential expression model (Lanz et al., 2019). Individual dataset reprocessing was performed using the *GEO2R* pipeline (Barrett et al., 2013); using methods mimicking Gemma’s gene expression processing as closely as possible to ensure consistency across datasets. The *GEOquery* package was used to extract data from Gene Expression Omnibus (v.2.66.0) (Barrett et al., 2013; Davis & Meltzer, 2007), and the gene expression data standardized through a log2 transformation and quantile normalization. Quality control included double-checking checking for extreme outlier samples by visualizing the top two principal components of variation in the data and verifying that log2 expression values fell within expected ranges (–5 to 12 for RNA-Seq, 4 to 15 for microarray). The *limma* pipeline (v.3.17) was used to quantify treatment-related differential expression for all genes included in the dataset (Ritchie et al., 2015; Smyth, 2005), using only the subset of samples relevant to the antipsychotic treatment vs. control frontal cortex analysis. The Benjamini-Hochberg method was used to correct nominal p-values for false discovery rate (FDR).

### Result extraction

Differential expression results for each of the eight antipsychotic treatment vs. control contrasts were extracted from either our individually reprocessed differential expression results (GSE66275, GSE4031.1) or the differential expression results provided by Gemma (GSE45229, GSE93918, GSE2547). Log2 Fold Changes (Log2FC) and corresponding t-statistics were extracted for each contrast. Standard errors (SE) and sampling variances (SV) were then calculated using those values (*full meta-analysis input:* **Table S1**). If more than one result represented a gene (e.g., multiple microarray probes targeting the same gene), the Log2FCs and SEs were averaged. Results extracted from the Gemma database were then aligned across datasets from the same species using EntrezID, whereas results extracted from the re-analyzed datasets were aligned by gene symbol. Results were further joined across species and annotations using a version of the Jackson Labs Mouse Ortholog Database (Baldarelli et al., 2024) that had been trimmed to one-to-one orthologs (Meta-Analysis Input: **Table S1**). A random-effects meta-analysis model was fit to the Log2FC and respective SV values for each gene using the *metafor* package (version 4.8-0; Viechtbauer, 2010). In order to be included within the meta-analysis, a gene was required to be present within at least 7 of the 8 treatment vs. control comparisons. The Benjamini-Hochberg method was used to correct nominal meta-analysis p-values for FDR using the *multtest* package (v.2.60.0; (Pollard, Dudoit, & van der Laan, 2005). Differentially expressed genes (DEGs) were defined using a 5% FDR threshold (FDR<0.05). Meta-analysis results for individual DEGs were visualized using forest plots, which illustrate the effect sizes (Log2FCs) and respective 95% confidence intervals from each study (the *forest.rma()* function from the *metafor* package).

### Assessment of Publication Bias, Result Heterogeneity, and Robustness

To evaluate the robustness and validity of the meta-analysis results, we conducted additional analyses to assess potential publication bias, residual heterogeneity, influential outliers, and the potential moderating factor of treatment type (first generation (typical) vs. second generation (atypical) antipsychotics). Potential publication bias was identified using Egger’s regression analysis (function *regtest()* in *metafor* v.4.8.0, (Viechtbauer, 2010), which detects funnel plot asymmetry. Egger p-values were corrected for FDR using the Benjamin-Hochberg method (*multtest* v.1.32.0, (Pollard, Dudoit, & van der Laan, 2005)) and genes with Egger FDR<0.05 were considered to show significant evidence of publication bias. Residual between-study heterogeneity was documented using common metrics extracted from the *metafor* meta-analysis object (Tau^2^ and associated SE, I^2^, H^2^) and evaluated using Cochran’s Q statistic (QE). After the QE nominal p-values were corrected for FDR using the Benjamin-Hochberg method (*multtest* v.1.32.0, (Pollard, Dudoit, & van der Laan, 2005)), genes with QE FDR<0.05 were considered to show significant heterogeneity across studies. To assess the influence of individual studies on meta-analysis results, the meta-analysis was repeated while iteratively excluding one study contrast at a time (*leave1out()* function in *metafor* (Viechtbauer, 2010) v.4.8.0). DEGs were considered to be overly dependent on individual studies if their significance dropped below a nominally significant p-value (p<0.05) upon removal of any single study.

To evaluate whether gene expression effects differed between first and second generation antipsychotics, we conducted a meta-regression analysis that included antipsychotic class as a co-variate, with general antipsychotic effects (intercept) centered between the two antipsychotic classes (*contrast.sum*: first-generation antipsychotics coded as −0.5, second-generation antipsychotics coded as 0.5). Significant effects of antipsychotic class were defined by having an FDR<0.05.

### Functional patterns

In order to identify potential functional gene pathways implicated in our results, fast Gene Set Enrichment Analysis (GSEA) was performed using the *fgsea* package (v.1.32.0; Korotkevich et al., 2021) to measure the enrichment of differential expression (upregulation or downregulation) in functional gene sets. A curated gene set database including both traditional gene ontology gene sets and brain-related gene sets was used (Brain.GMT: (M. H. Hagenauer et al., 2024; Subramanian et al., 2005)). Enrichment of differential expression within each gene set was calculated by applying *fgsea* to the meta-analysis results ranked by estimated treatment effect size (Log2FC). The analysis was conducted using 10,000 permutations, with gene sets restricted to those containing between 10 and 1000 genes. In instances where multiple Log2FCs mapped to the same gene symbol, effect sizes were aggregated by computing the mean prior to ranking. A second “non-directional” fGSEA analysis was conducted using the absolute values of each gene’s Log2FC. Gene sets with a significant enrichment of differential expression (FDR<0.05) in either the directional or non-directional analysis were then examined to determine whether the leading genes driving the enrichment included DEGs identified by the meta-analysis.

### Exploratory: Cross-species comparisons

The interpretation of our planned meta-analysis was limited by the fact that the only rodent studies available focused on male subjects and treatment protocols that lasted for less than a month. Moreover, rodents have an anatomical organization in the frontal cortex that is distinct from primates and humans (Preuss & Wise, 2022). To provide a greater cross-species perspective, we compared the rodent meta-analysis results to previously published differential expression results from two primate studies. One study used RNA-Seq to profile the dorsolateral PFC (DLPFC) of healthy male and female rhesus macaques treated with either clozapine (5.2 mg/kg daily), high dose haloperidol (4 mg/kg daily), low dose haloperidol (0.14 mg/kg daily), or placebo for six months (Schulmann et al., 2023). The full differential expression results were reported in the supplement (16,080 genes). The second study (Martin et al., 2015) used two different human microarray platforms (Affymetrix HU6800 and HG-U95A/HG-U95Av2) to profile the PFC of crab-eating macaques that were either controls (vehicle-treated daily) or modeling psychosis by daily treatment with PCP (1 mg/kg) for 14 days. The control animals were then treated with haloperidol (1.5 mg/kg daily), olanzapine (0.75 mg/kg daily), or vehicle (daily) for 28 days, while the psychosis animals were either treated with olanzapine (0.75 mg/kg daily) or vehicle (daily) while continuing PCP (2 mg/kg for 28 days) (**Table 1**). This study also included acute haloperidol and olanzapine treatment groups which were not used in our analyses. The full differential expression results from both microarray platforms were reported in the supplement.We filtered out results from probes that mapped to more than one gene symbol, leaving 5942 probes representing 5176 gene symbols (HU6800), and 7224 probes representing 5175 gene symbols (HG-U95A/HG-U95Av2).

To make initial comparisons between our rodent results and the rhesus macaque findings (Schulmann et al., 2023), we averaged the log2FCs across macaque clozapine, low-dose haloperidol, and high-dose haloperidol treatment groups, and calculated the minimum nominal p-value and FDR across treatment groups. For comparisons with the crab-eating macaque findings (Martin et al., 2015), the gene expression ratios for the chronic antipsychotic treatment vs. vehicle comparisons (haloperidol vs. vehicle, olanzapine vs. vehicle, olanzapine+PCP vs. PCP) were extracted from each of the platforms and log(2) transformed to produce log2FCs. In the case of the Olanzapine+PCP vs. PCP comparison, the direction of effect was originally reported using a reversed definition (PCP as treatment vs. Olanzapine+PCP as reference) within the results from one platform (HG-U95A/HG-U95Av2), so it was necessary to invert the ratio (1/ratio) before performing log2 transformation. Log2FCs for probes representing the same gene symbols were averaged within each platform, and then the log2FCs for each gene were averaged across platforms and antipsychotic treatment conditions. The minimum nominal p-value and FDR were also calculated across probes representing the same gene symbols within each platform, and then for each gene across platforms and antipsychotic treatment conditions.

To align our rodent results with the rhesus macaque findings (Schulmann et al., 2023), we initially identified reliable one-to-one orthologs across rodents and rhesus macaques for each of our DEGs by mapping between mouse Entrez Gene IDs to macaque Ensembl gene IDs using bioDBnet (accessed June 25, 2025; referencing EntrezGene release date June 20, 2025, updated June 22, 2025 and Ensembl 113 release date May 13, 2025, updated May 18, 2025) (Mudunuri et al., 2009). For more in depth comparisons, we leveraged the human orthologs provided by the original publication for each of the macaque genes (Schulmann et al., 2023). We then used the HGNC Comparison of Orthology Predictions (HCOP) database (downloaded from https://www.genenames.org/tools/hcop/ on Aug 21, 2025) to identify the human orthologs for all of the mouse genes included in our meta-analysis results (downloaded file: “human_mouse_hcop_fifteen_column.txt”). We selected mouse-human gene orthologs that either 1) used the same official gene symbol, or 2) were supported by the largest number of databases referenced by HCOP, with ties broken by row index. To compare our rodent results to the crab-eating macaque findings (Martin et al., 2015), we similarly aligned results using the human gene symbols provided in the original publication, as the results were derived from human microarray platforms.

### Exploratory: Meta-analysis incorporating rodent and non-human primate data

To increase the power and generalizability of our meta-analysis findings, we decided to run an exploratory meta-analysis study that leveraged both the rodent antipsychotic studies and aforementioned primate studies. Unfortunately, we were unable to obtain either the raw data or full statistical reporting needed to run an effect size meta-analysis using the crab-eating macaque study (Martin et al., 2015). Therefore our exploratory meta-analysis only incorporated the rhesus macaque results (Schulmann et al., 2023). From the rhesus macaque study, we extracted the antipsychotic effect sizes (Log2FCs) and associated sampling variance for each of the treatment comparisons (clozapine (5.2 mg/kg) vs. placebo, high dose haloperidol (4 mg/kg) vs placebo, low dose haloperidol (0.14 mg/kg) vs. placebo) for each of the macaque genes that had mouse orthologs. We then added these differential expression results to our previous meta-analysis using the standardized pipeline overviewed above. In order to be included within the meta-analysis, we required that a gene be present within at least 7 of the 11 treatment vs. control comparisons. This meant that all of the genes included in our previous planned rodent meta-analysis would be represented in the exploratory meta-analysis, however, it is worth noting that some genes (2,504, or 19%) in the exploratory meta-analysis still had results that were exclusively derived from the rodent studies.

The findings from the exploratory meta-analysis are likely to have greater power and generalizability than the findings from our planned rodent meta-analysis, but there is an enhanced possibility for false discovery in exploratory (vs. planned) analyses. Therefore, within our results, we decided to emphasize findings that were significant in both our original planned meta-analysis and that remained significant following the addition of the rhesus macaque data in our exploratory meta-analysis.

### Exploratory: Comparisons with human clinical data

To explore the clinical relevance of our findings, we compared our meta-analysis results to the results from several relevant differential expression analyses from human patients. One analysis examined the effects of schizophrenia in the DLPFC of individuals who did not have antipsychotics detectable at time of death by post-mortem toxicology (Antipsychotic-, n=23), or who did have antipsychotics detectable by post-mortem toxicology (Antipsychotic+ n=65), subgrouped by treatment class: atypical, typical, or mixed (Schulmann et al., 2023) (**Table 1**). To simplify comparisons, we calculated the average effect of schizophrenia (Log2FC) across all Antipsychotic+ groups, and then compared them to the effect of schizophrenia (Log2FC) within the Antipsychotic-group. We also compared our findings to two large-scale meta-analyses of human post-mortem transcriptomic data (Gandal et al., 2018). We extracted the differential expression results for the schizophrenia vs. control comparisons in the microarray (cortex) and RNA-seq (DLPFC) meta-analyses from supplementary *Tables S1* and *S2*, respectively (**Table 1**). These differential expression results were merged by human Ensembl gene ID and joined to the rodent-macaque dataframe to allow comparisons.

## Results

### Planned Rodent Meta-Analysis

We used systematic inclusion and exclusion criteria to identify datasets for our rodent antipsychotic meta-analysis. Five datasets met our criteria, containing a total of eight Antipsychotic vs. Control comparisons and a modest collective sample size of *n*=68 subjects (**Table 1**). The treatments included in the meta-analysis encompassed both typical (*first generation:* haloperidol, collective *n*=16) and atypical (*second generation:* quetiapine, clozapine, olanzapine, risperidone, collective *n*=26) antipsychotics, and normally lasted between 21-28 days. Subjects included both adult mice and rats, which, to our knowledge, were all male. Gene expression was predominantly measured using microarray (Affymetrix platform), with only one dataset using RNA-Seq.

In order to be included within the meta-analysis, we required that a gene be represented in at least 7 of the 8 antipsychotic vs. control comparisons. Of the eligible 12,215 genes, 12,190 genes produced stable meta-analysis estimates (*full results:* **Table S2**). In total, 63 genes were significantly differentially expressed following antipsychotic treatment (**Figure 2**: “DEG”: FDR<0.05), 45 of which were upregulated (**Table S3**) and 18 downregulated (**Table S4**).

**Figure 2.**
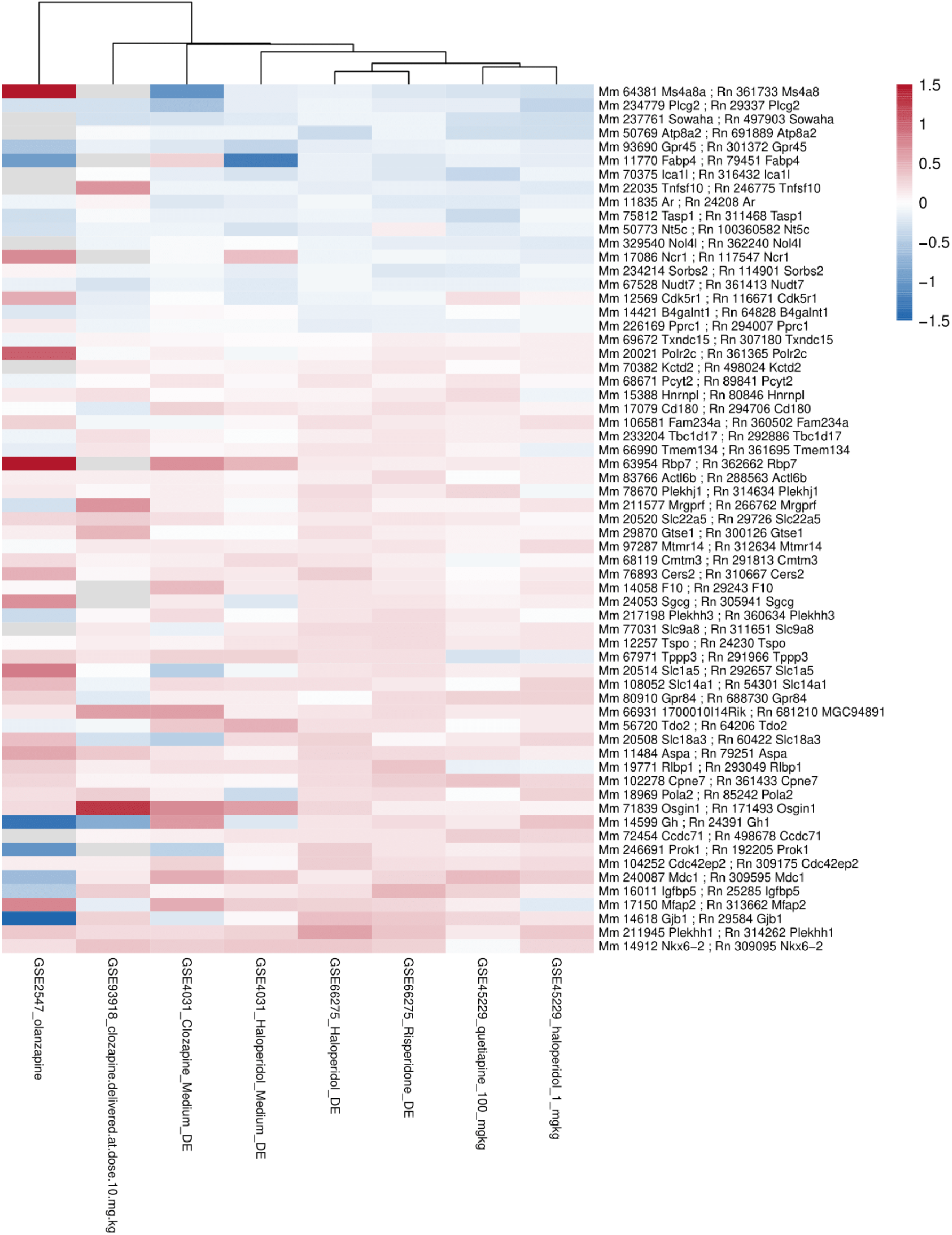
A hierarchically-clustered heatmap illustrating the antipsychotic vs. control effect sizes across all group comparisons in the rodent meta-analysis for each of the top differentially expressed genes (DEGs). The heatmap displays the antipsychotic vs. control effect sizes (Log(2) fold changes or Log2FCs) for each of the 63 differentially expressed genes identified through the meta-analysis as significantly differentially expressed (FDR<0.05). Results are scaled and color-coded by log2FC, ranging from −1.5 to +1.5 (blue to red; downregulation to upregulation). Rows represent individual genes, annotated with both mouse (Mm) and rat (Rn) identifiers and gene symbols, and each column corresponds to a specific treatment vs. control comparison, identified by Gene Expression Omnibus dataset id (GSE…) and treatment used. Hierarchical clustering is applied and used to group columns with similar expression profiles.

### Rodent Meta-Analysis: Assessment of Result Robustness and Validity

Amongst our top meta-analysis results (63 DEGs), we did not see evidence of clustering by antipsychotic class, species, dissection, or platform (**Figure 2**). None of the 63 DEGs showed significant between-study heterogeneity (QE FDR<0.05), and only one gene showed weaker, nominal evidence of heterogeneity (QE p<0.05). Similarly, when antipsychotic class was included as a co-variate in an exploratory meta-regression analysis, the only gene that showed a significant effect of antipsychotic class (first generation vs. second generation: FDR<0.05) was *Neuromedin U Receptor 1* (*Nmur1*), which was not one of the antipsychotic DEGs identified in our original-analysis (*full results:* **Table S5**). Although our power is limited, these findings are consistent with the conclusion that the DEGs identified by our planned antipsychotic meta-analysis are generally responsive to both classes of antipsychotics.

Our rodent meta-analysis results appeared both robust and valid by many other metrics. Overall, we had deemed the risk of systematic bias low for any particular gene-level result, as the original publications conducted full genome analyses and the datasets were re-analyzed using standardized pipelines. However, bias in the literature against negative results might inflate effect sizes represented in publicly-released datasets, so we further assessed potential publication bias using Egger’s regression analysis. This analysis identified 39 genes with significant bias after FDR correction (Egger FDR<0.05, **Table S6**), which did not include any of the 63 antipsychotic DEGs identified by our meta-analysis. Four of our 63 DEGs showed weaker, nominal evidence of potential publication bias (Egger p<0.05: *Ccdc71*, *Gjb1*, *Cdk5r1*, and *Tnfsf10*). Iterative study-removal analysis demonstrated that only 5 of our 63 DEGs dropped below nominal significance (p>0.05) when a single contrast was removed (*Mfap2, Tppp3, Gtse1, Gh*, and *Cdk5r1*), indicating that our findings are also robust.

### Exploratory Rodent & Macaque Meta-Analysis

The 63 DEGs that we identified in our planned meta-analysis were identified in rodents, which have an anatomical organization in the frontal cortex that is distinct from primates and humans (Preuss & Wise, 2022). These studies were also predominantly focused on male subjects and treatment protocols that lasted for less than a month. To increase the power and generalizability of our results, we performed a meta-analysis that included both the aforementioned rodent datasets and RNA-Seq data from the DLPFC of rhesus macaques that had been treated for six months with clozapine, high dose haloperidol, low dose haloperidol, or placebo (**Table 1**). By including the rhesus macaque data, our total sample size increased to *n*=101, with 11 total antipsychotic vs. control comparisons. Similar to the planned meta-analysis, we required that a gene be represented in at least 7 of the 8 comparisons to be included in the exploratory meta-analysis. Of the eligible 13,121 genes, 10,617 now also included macaque data in the meta-analysis, and 13,107 produced stable meta-analysis estimates (*full results*: **Table S7**). In total, 117 genes were differentially expressed following antipsychotic treatment (**Table S8 & S9**, DEGs: FDR<0.05). Of these, 31 had been identified as differentially expressed in our original planned meta-analysis, 16 of which now included macaque data in the estimate (listed in **Table S3,** example forest plots: **Figure 3**).

**Figure 3.**
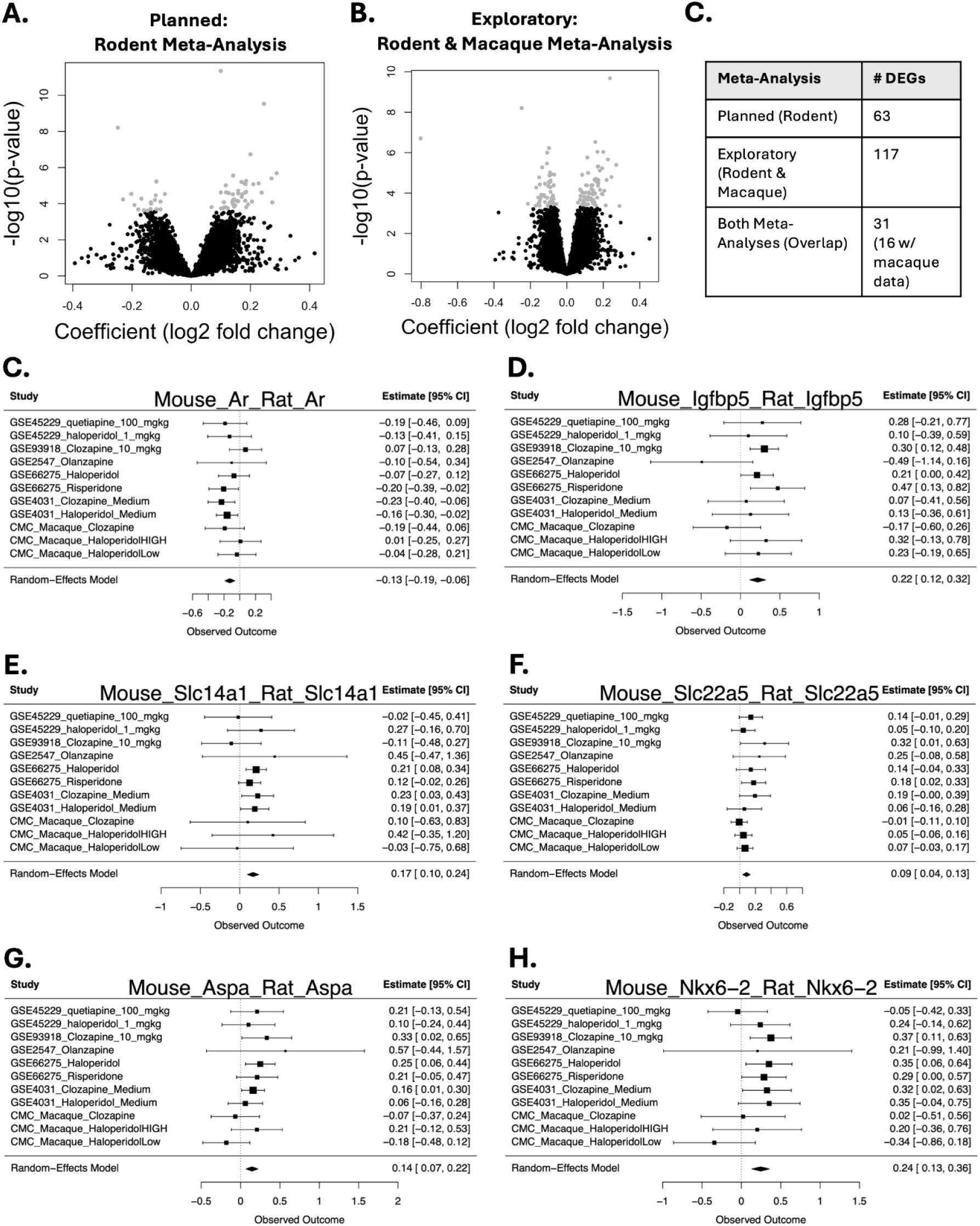
Examples of genes that were differentially expressed following antipsychotic treatment in both our planned rodent meta-analysis and larger, exploratory meta-analysis that included both macaques and rodents. **A-B.** Volcano plots illustrating differential gene expression in the planned rodent meta-analysis (**A**) and exploratory rodent and macaque meta-analysis (**B**). Volcano plots depict the relationship between effect size (log(2) fold change or Log2FC) and statistical significance. Points highlighted in grey signify significantly differentially expressed genes (DEGs: FDR<0.05). Negative Log2FC indicates downregulated genes following antipsychotic treatment and positive Log2FC indicates upregulated genes. **C.** A table showing the number of significant DEGs (FDR<0.05) identified within the planned rodent meta-analysis and larger, exploratory rodent and macaque meta-analysis, as well as their overlap. Please note that the larger, exploratory rodent and macaque meta-analysis included results from some genes that lacked macaque data, as long as there were at least seven antipsychotic vs. control comparisons available within the rodent data. **C-H.** Example forest plots from genes that were differentially expressed within the planned rodent meta-analysis and larger, exploratory rodent and macaque meta-analysis. Rows illustrate the effect of antipsychotic medication on gene expression in the frontal cortex and its Log2FC (squares) with 95% confidence intervals (whiskers) for each of the datasets and the meta-analysis random effects model. Forest plots allow for visual inspection of the consistency and magnitude of effects across the datasets (indicated with their Gene Expression Omnibus accession number or consortium (CommonMind: CMC) and the specific antipsychotic treatment used in the comparison). **C.** Antipsychotics produced a down-regulation of Androgen receptor (Ar), which encodes a nuclear receptor that binds testosterone and dihydrotestosterone, **D.** Antipsychotics produced an upregulation of Insulin growth factor binding protein 5 (Igfbp5), which encodes a protein that enables insulin-like growth factor I binding activity, **E.** Antipsychotics produced an upregulation of Solute carrier family 14 member 1 (Slc14a1), which encodes a urea transporter protein**, F.** Antipsychotics produced an upregulation of Solute carrier family 22 member 5 (Slc22a5), which encodes a sodium-ion dependent, high affinity carnitine transporter, **G.** Antipsychotics produced an upregulation of Aspartoacylase (Aspa), which encodes an enzyme that catalyzes the conversion of N-acetyl-L-aspartic acid (NAA) to aspartate and acetate that is thought to help maintain white matter. **H.** Antipsychotics produced an upregulation of NK6 Homeobox 2 (Nkx6-2), which encodes a transcription factor that is thought to be important for myelination.

Despite both species differences and protocol differences between the macaque and rodent studies, the hierarchically-clustered heatmap for the top DEGs did not reveal clear clustering by species (**Fig S1**). Similarly, none of the top DEGs showed significant between-study heterogeneity (QE FDR<0.05) and only two showed nominal between-study heterogeneity (QE p<0.05). Therefore, the antipsychotic DEGs identified in the exploratory meta-analysis are likely to be generalizable across species and treatment conditions.

The exploratory results also appeared generally robust and valid, with only three DEGs dropping below nominal significance (p<0.05) when a single contrast was removed (*Gtse1, Gh, Cdhr5*), and none showing significant evidence of potential publication bias (Egger FDR<0.05, **Table S7**). However, the results of exploratory analyses are more likely to reflect false discovery than those of planned comparisons. Therefore, we decided to focus our remaining discussion on findings that were significant in our original planned meta-analysis and that remained significant following the addition of the rhesus macaque data in our exploratory meta-analysis.

### Functional patterns

Functional patterns within the meta-analysis results were examined using fast Gene Set Enrichment Analysis (*fGSEA*) using a database containing both traditional gene ontology gene sets and brain-related gene sets (Brain.GMT). Within the planned rodent meta-analysis, 14 gene sets (out of 10,147) were significantly enriched (FDR<0.05) with down-regulation following antipsychotic treatment, and 50 enriched with upregulation. A non-directional version of the analysis identified 170 gene sets enriched with differential expression following antipsychotic treatment (FDR<0.05), 34 of which were also significant in the directional version of the analysis. Across these findings, visible patterns of functional convergence were observed (**Table 2**, *full results:* **Tables S10 & S11**). Upregulated gene sets were often related to glial cell types or glial development, with a particularly large number of upregulated gene sets (14) related to oligodendrocyte development and myelination. There were also upregulated gene sets related to astrocytes and radial glia, as well as fibroblast-like cell pathways. Other cell types represented in the gene set enrichment results included microglia, progenitor cells, neurogenesis, and ependymal cells. Unique to the non-directional results were a large number of gene sets (20) related to vascular cell types (endothelial cells, pericytes), the cardiovascular and circulatory system, and blood pressure. Notably, many of the gene sets enriched with differential expression (17) - both up and down-regulation - were related to fear conditioning or stress responses, as well as stress-related disorders (Major Depressive Disorder). Other themes included growth and development, molecular transport, cellular signalling (hormone, G-protein, enzyme-linked), and metabolic processes/symptoms.

**Table 2.**
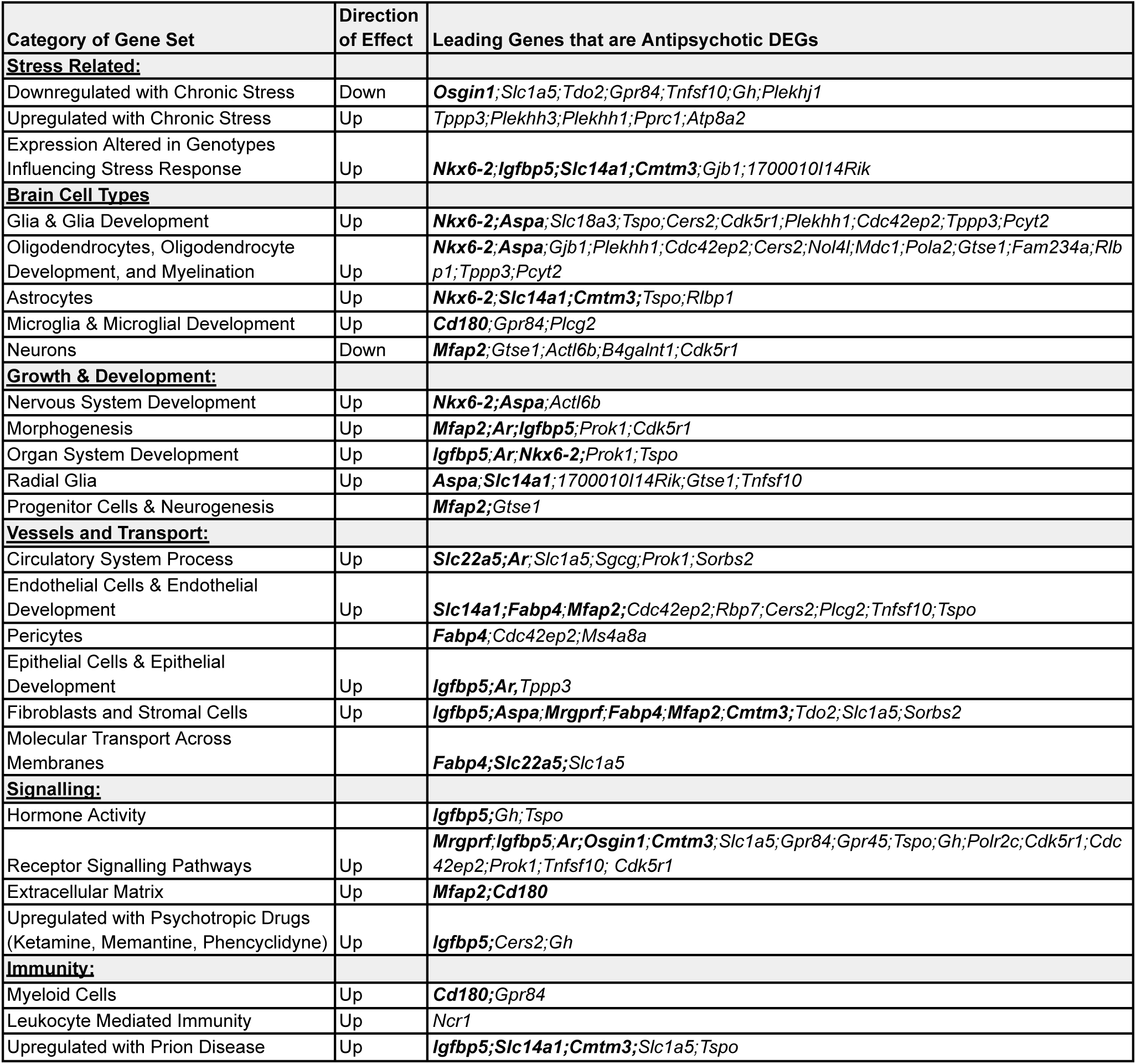
An overview of functional gene sets containing antipsychotic DEGs that were enriched with antipsychotic-related differential expression. The table overviews categories of functional gene sets that showed significant enrichment (FDR<0.05) with antipsychotic-related differential expression within the original planned rodent meta-analysis as well as in the larger, exploratory rodent & macaque meta-analysis. For conciseness, the table only overviews functional categories that were represented by more than one gene set and that contained antipsychotic DEGs. If the category contained gene sets that were significant within a directional analysis (for either the planned or exploratory meta-analysis), the direction of effect is provided, with “Up” signifying an enrichment with upregulation and “Down” signifying an enrichment with downregulation. The leading genes that were antipsychotic DEGs within our planned meta-analysis are listed for each category by official mouse gene symbol, with bold text indicating that the DEG also remained significant in the larger, exploratory rodent & macaque meta-analysis and had rhesus macaque data.

These patterns were largely replicated when performing a similar fGSEA analysis using the exploratory rodent & macaque meta-analysis results. This analysis revealed a larger number of gene sets that were significantly enriched with differential expression (FDR<0.05, *full results:* **Tables S12 & S13**), with 235 (out of 10,417) gene sets identified as enriched in the directional version of the analysis and 233 gene sets identified as enriched in the non-directional version, but the identified gene sets were thematically similar to the patterns identified in the planned rodent meta-analysis results. The most noteworthy difference was a large increase in the number of enriched gene sets related to leukocytes and immunity, which all showed upregulation. Some gene sets that were previously only enriched in the non-directional version of the analysis were now also significant in the directional version of the analysis (*e.g.,* the upregulation of cardiovascular and blood vessel related gene sets). Notably, there was also upregulation within a set of genes previously identified as upregulated in the cortex of individuals with schizophrenia in a large meta-analysis (Gandal et al., 2018), led by the DEGs *Adenosine A2b receptor* (*Adora2b*), which encodes a g-protein coupled adenosine receptor, and *Solute carrier family 14 member 1* (*Slc14a1*), which encodes a urea transporter.

### Exploratory: Overlap with Human Schizophrenia Results

To explore the potential clinical relevance of our findings, we decided to compare our meta-analysis results to 1) The effects of schizophrenia within the DLPFC of subjects that were on antipsychotics (antipsychotic+) vs. not on antipsychotics (antipsychotic-) at the time of death (Schulmann et al., 2023), 2) The effects of schizophrenia on the frontal cortex within larger meta-analyses, with treatment status unknown (Gandal et al., 2018). Of particular interest would be genes that show consistent antipsychotic effects in animals that are *a reversal* of the effects of psychotic disorders, especially a reversal of diagnosis effects observed in subjects that were not on antipsychotics at the time of death (*i.e.,* effects that might be therapeutic). Another point of interest would be genes that show consistent antipsychotic effects in animals that *resemble* the effects of psychotic disorders, especially in subjects that were on antipsychotics at the time of death (*i.e.,* effects of antipsychotics that might be mistaken for disease processes).

We narrowed our focus to genes with the most consistent antipsychotic effects in animals, as defined as the 31 antipsychotic DEGs identified in the planned rodent meta-analysis that remained significant in the larger, exploratory rodent & macaque meta-analysis (**Table 3**). Twelve of these 31 genes also had results available from crab-eating rhesus macaques (Martin et al., 2015), which typically showed weak congruent Log2FCs (9 of 12 genes), providing additional support. Within the human schizophrenia datasets, there was data available for 23 of the 31 genes. Several of the antipsychotic DEGs had either significant (FDR<0.05) or multiple nominal (p<0.05) effects of schizophrenia within the post-mortem frontal cortex: *Insulin growth factor binding protein 5 (Igfbp5),* which encodes a protein that enables insulin-like growth factor I binding activity, *Taspase 1 (Tasp1)*, which encodes a protease, and multiple members of the solute carrier family (SLC) — *Solute Carrier Family 14 Member 1* (*Slc14a1*), *Solute Carrier Family 1 Member 5* (*Slc1a5*), and *Solute Carrier Family 22 Member 5* (*Slc22a5*) — which encode transporters implicated in urea transport, glutamine/glutamate transport, and high affinity carnitine transport, respectively. All of these genes showed similar differential expression in individuals with schizophrenia as what was observed in response to antipsychotics in animals, implying that the observed diagnosis-related differential expression may represent treatment effects.

**Table 3.** Exploratory: Antipsychotic treatment effects within primates and human post-mortem schizophrenia datasets. This table presents genes that were consistently differentially expressed (DEGs FDR<0.05) following antipsychotic treatment both in our planned rodent meta-analysis and larger, exploratory rodent & macaque meta-analysis in relationship to their corresponding differential expression findings in human post-mortem schizophrenia datasets. Antipsychotic DEGs that lacked representation in both the primate and human datasets are not shown (Ms4a8a, Gpr84, 1700010I14Rik, Prok1). The “Antipsychotic+ Rodent Meta-Analysis Log2FC” column shows the log2 fold change of antipsychotics vs. control within the planned rodent meta-analysis. The “Antipsychotic+ Rhesus Macaque AVE_Log2FC” column represents the average log2 fold change across macaque clozapine, low-dose haloperidol, and high-dose haloperidol treatment groups (Schulmann et al., 2023). The “Antipsychotic+ Crab-eating Macaque AVE_Log2FC” column represents the average log2 fold change across macaque haloperidol and olanzapine treatment groups (in controls and the psychosis model). The “Antipsychotic+ SCHIZ AVE_Log2FC” column represents the average diagnosis log2 fold change across the schizophrenia subjects on atypical, typical, and mixed antipsychotic treatments, as detected by post-mortem toxicology (Schulmann et al., 2023), and the “Antipsychotic-SCHIZ Log2FC” column represents the diagnosis log2 fold change for the schizophrenia subjects that were not on antipsychotic medication at the time of death (Schulmann et al., 2023). The “SCHIZ Gandal RNA-Seq Log2FC” and “SCHIZ Gandal Microarray Log2FC”columns show the log2 fold changes for schizophrenia from large RNA-Seq and microarray meta-analyses, with treatment status unknown (Gandal et al., 2018). The color scale indicates the magnitude and direction of gene expression (red = upregulation, blue = downregulation), with bold text indicating a nominal association (p<0.05) in any of the referenced comparisons, and underline indicating a significant association (FDR<0.05).

| Mouse Symbol | Antipsychotic+<br>Rodent<br>Meta-Analysis<br>Log2FC | Antipsychotic+<br>Rhesus<br>Macaque<br>AVE_Log2FC | Antipsychotic+<br>Crab-Eating<br>Macaque<br>AVE_Log2FC | Antipsychotic+<br>SCHIZ<br>AVE_Log2FC | Antipsychotic-<br>SCHIZ<br>Log2FC | SCHIZ<br>Gandal<br>RNA-Seq<br>Log2FC | SCHIZ<br>Gandal<br>Microarray<br>Log2FC |
| --- | --- | --- | --- | --- | --- | --- | --- |
| *Fabp4* | -0.17 | -0.22 | 0.02 | NA | NA | NA | 0.00 |
| *Ar* | -0.14 | -0.07 | 0.04 | 0.08 | 0.06 | 0.00 | 0.00 |
| *Tasp1* | -0.14 | -0.05 | NA | -0.03 | -0.03 | -0.06 | -0.09 |
| *Ncr1* | -0.12 | NA | NA | NA | NA | NA | 0.03 |
| *Nudt7* | -0.12 | 0.00 | NA | 0.05 | -0.08 | -0.07 | NA |
| *Cd180* | 0.11 | 0.10 | 0.03 | NA | NA | NA | 0.01 |
| *Tmem134* | 0.12 | 0.05 | NA | -0.05 | -0.05 | 0.00 | 0.01 |
| *Rbp7* | 0.12 | NA | NA | 0.06 | 0.18 | 0.04 | NA |
| *Mrgprf* | 0.14 | 0.16 | NA | NA | NA | NA | NA |
| *Slc22a5* | 0.14 | 0.04 | NA | 0.06 | -0.02 | 0.07 | 0.07 |
| *Gtse1* | 0.14 | NA | NA | NA | NA | NA | 0.03 |
| *Cmtm3* | 0.14 | 0.04 | NA | 0.06 | 0.03 | 0.08 | NA |
| *F10* | 0.15 | 0.21 | 0.03 | NA | NA | NA | -0.04 |
| *Slc1a5* | 0.16 | NA | 0.02 | 0.25 | 0.20 | 0.28 | 0.01 |
| *Slc14a1* | 0.17 | 0.16 | -0.01 | 0.36 | 0.60 | 0.39 | 0.46 |
| *Tdo2* | 0.18 | NA | 0.01 | NA | NA | 0.08 | -0.03 |
| *Slc18a3* | 0.18 | NA | 0.03 | NA | NA | NA | 0.01 |
| *Aspa* | 0.18 | -0.01 | 0.02 | -0.12 | -0.21 | -0.01 | 0.03 |
| *Osgin1* | 0.19 | 0.07 | NA | -0.06 | -0.06 | -0.04 | -0.03 |
| *Gh* | 0.19 | NA | NA | NA | NA | NA | 0.02 |
| *Ccdc71* | 0.20 | 0.09 | NA | -0.01 | 0.02 | 0.02 | NA |
| *Igfbp5* | 0.24 | 0.13 | 0.03 | 0.20 | 0.05 | 0.14 | 0.06 |
| *Mfap2* | 0.25 | 0.15 | 0.06 | NA | NA | NA | NA |
| *Gjb1* | 0.27 | NA | 0.04 | -0.15 | -0.04 | -0.11 | -0.05 |
| *Nkx6-2* | 0.29 | -0.04 | NA | -0.05 | 0.17 | 0.12 | NA |

However, when examining results from the frontal cortex of subjects with schizophrenia that were *not* on antipsychotics at the time of death (Schulmann et al., 2023), differential expression was often similar to that observed in the other diagnosis comparisons, although the effects within this smaller subgroup typically did not meet a threshold for significance (**Table 3**). The only exception was *Slc22a5,* which showed upregulation in all schizophrenia analyses except within the antipsychotic negative schizophrenia subgroup, which showed weak down-regulation. Altogether, these findings suggest that antipsychotic treatment may account for some effects within schizophrenia post-mortem datasets, but that these effects are more likely to be driven by chronic treatment exposure and cannot be disambiguated from disease processes using evidence of treatment at time of death via post-mortem toxicology.

The most convincing evidence for antipsychotics reversing diagnosis effects came from *Aspartoacylase (Aspa),* which encodes an enzyme that catalyzes the conversion of N-acetyl-L-aspartic acid (NAA) to aspartate and acetate that is thought to help maintain white matter. *Aspa* was down-regulated in subjects with schizophrenia that tested negative for antipsychotics at time of death (Log2FC=-0.21), but upregulated by antipsychotic treatment. However, the down-regulation observed in the Antipsychotic-subjects was not significant, potentially due to low subgroup sample size.

## Discussion

Psychotic disorders represent one of the most expensive and straining classes of psychiatric disorders, necessitating proper, targeted, and effective treatment strategies (Christensen et al., 2020). Antipsychotic medications serve as the primary pharmacological therapy; however, the effects of antipsychotic treatment on the brain are not fully understood. Exploring effects within the frontal cortex is critical as it plays a significant role in cognitive executive functions such as decision-making, emotional regulation, social behavior, and analytical thinking (D. T. Jones & Graff-Radford, 2021). Psychotic disorders are known to cause significant structural and functional changes within the frontal cortex, all of which may influence the body’s executive functions (Bilecki & Maćkowiak, 2023).

In order to determine the effects of antipsychotic treatment on the frontal cortex, we conducted a systematic meta-analysis of publicly available rodent transcriptional profiling datasets. We utilized a selection of five public datasets (GSE45229, GSE93918, GSE2547, GSE4031.1, and GSE66275), providing us with eight drug vs. control comparisons that were derived from both atypical and typical psychotic drug classes. The statistical power was modest, with a final collective sample size of *n*=68. Altogether, we obtained stable meta-analysis estimates for 12,190 genes, with 63 genes being significantly differentially expressed (FDR<0.05), most of which (45/63) were upregulated. This antipsychotic differential expression was enriched within gene sets showing visible patterns of functional convergence with respect to oligodendrocyte development, vasculature, and stress responses.

To improve the generalizability of our findings across species, sexes, and treatment protocols, we ran a second, exploratory meta-analysis that included macaque as well as rodent data, increasing our collective sample size to *n*=101. This exploratory meta-analysis expanded upon and confirmed many of the findings from our planned rodent meta-analysis. However, exploratory analyses have a greater risk of false discovery, therefore, we have highlighted findings in our discussion that were both significant in our original planned meta-analysis and remained significant in the exploratory meta-analysis following the addition of the rhesus macaque data.

### Differential Expression Related to Previously Implicated Neurotransmitter Pathways

Many theories of schizophrenia have focused on the role of dopamine in the generation of psychosis (Stahl, 2018). Both first generation (typical) and second generation (atypical) antipsychotics target dopaminergic function, especially dopamine D2 receptors (Julayanont & Suryadevara, 2021; H. Y. Meltzer, 2013; Siafis et al., 2023). Therefore, it was surprising that genes associated with dopamine function were not highlighted amongst our top results. Instead, our top findings included the upregulation of two genes with clear roles in neurotransmitter systems targeted by newer therapeutics: *Tryptophan 2,3-Dioxygenase (Tdo2),* which encodes an enzyme that plays a critical rate-limiting step in the metabolism of the serotonin precursor tryptophan, and *Solute carrier family 18 member A3* (*Slc18a3)*, which encodes the Vesicular Acetylcholine Transporter (VAChT) responsible for packaging acetylcholine into synaptic vesicles.

The upregulation of *Tdo2* in response to antipsychotics could play a therapeutic role by directing the metabolism of tryptophan away from the serotonin pathway, reducing overall serotonin levels. Recent evidence suggests that serotonin release is elevated within the frontal cortex of individuals with schizophrenia, particularly in association with negative symptoms (Osugo et al., 2026). Atypical antipsychotics modulate serotonergic as well as dopaminergic neurotransmission, including antagonizing serotonin 5-HT_2_A receptors (H. Y. Meltzer, 2013). Serotonergic signaling is also known to be strongly interconnected with glutamatergic and dopaminergic neural networks implicated in psychosis (Stahl, 2018). Within mice, mutations in *Tdo2* have been associated with altered prepulse inhibition (Baldarelli et al., 2024; Wilson et al., 2026), which is an operational measure of sensorimotor gating (W. Li et al., 2021; San-Martin et al., 2020). These findings suggest that directly targeting *Tdo2* can modulate psychosis-like behavior in animal models (W. Li et al., 2021; San-Martin et al., 2020).

The upregulation of *Slc18a3* in response to antipsychotics may also play a therapeutic role by increasing the amount of acetylcholine available for synaptic release. Acetylcholine is implicated within cognitive processes including attention, learning, and memory - domains that are frequently impaired in schizophrenia. Cholinergic signaling interacts with dopaminergic neurotransmission, and alterations in cholinergic function may influence psychotic symptoms and cognitive dysfunction through modulation of dopamine release (Saint-Georges et al., 2025). Recently, Cobenfy (xanomeline and trospium chloride), which primarily activates M1 and M4 muscarinic acetylcholine receptors, was approved for the treatment of schizophrenia (Commissioner, 2024), following many decades of research documenting cholinergic disruption in schizophrenia (Saint-Georges et al., 2025) and the utility of cholinergic agonists as antipsychotics (Yohn et al., 2022). Our findings suggest that VAChT may also be a potentially useful target, and suggest that previous PET evidence linking increased VAChT binding to schizophrenia symptom severity may reflect antipsychotic usage (Weinstein et al., 2024).

However, although intriguing, both the *Tdo2* and *Slc18a3* findings should be interpreted as preliminary, as they were significant within both our planned and exploratory meta-analyses, but based exclusively on rodent data. Findings from the rhesus macaques were absent, and findings from crab-eating macaques and human patients, although directionally congruent, were relatively weak, requiring future validation.

### Differential Expression Related to Oligodendrocytes

Previous research has established a clear link between antipsychotic treatment for psychotic disorders to oligodendrocytes and white matter, particularly in the PFC. Oligodendrocytes are primarily responsible for myelin sheath formation and development in the central nervous system. Schizophrenia has been consistently correlated with oligodendrocyte dysfunction and myelin abnormalities as well as altered expression of myelin-related genes (e.g., *OLIG2, CNP, and NRG1*) (Gouvêa-Junqueira et al., 2020). These disruptions in oligodendrocyte function have been associated with the disordered executive function and development commonly associated with schizophrenia and other psychotic illnesses (Gouvêa-Junqueira et al., 2020; Mighdoll et al., 2015). Additionally, white matter, the brain tissue primarily consisting of myelinated axons, serves as an essential component of distributed neural networks, critical for the structural basis of human behavior (Crocker & Tibbo, 2018). Psychotic disorders are associated with reductions in white matter development and structure, correlating with poor cognitive function and neural disconnectivity (Crocker & Tibbo, 2018; Peters & Karlsgodt, 2015).

Our meta-analysis findings reinforced the potential efficacy of antipsychotics on schizophrenia-related white matter and oligodendrocyte deficits, as 14 of 50 upregulated pathways in our planned meta-analysis were related to oligodendrocyte development or oligodendrocyte-related function. This pattern was confirmed in our larger, exploratory rodent-macaque meta-analysis. Top DEGs that were leading genes within these enriched upregulated pathways, such as *Nkx6-2 (NK6 Homeobox 2), Aspa (Aspartoacylase)*, and *Gap Junction Protein Beta 1 (Gjb1),* are known for their role in oligodendrocyte function and myelination, and may reflect potential for remyelination or prevention of further demyelination (Chelban et al., 2017; Franklin & Simons, 2022). Myelin-related gene expression was also previously noted to be decreased in crab-eating macaques modeling psychosis (PCP treated) in a manner that was reversed by antipsychotics (Martin et al., 2015). Moreover, mutations in *Aspa* have been associated with decreased prepulse inhibition in mice (Baldarelli et al., 2024; Wilson et al., 2026), implying that the relationship between myelin-related gene expression and psychosis-like behavior may be causal. Although not significant, *Aspa* was also decreased in the DLPFC of individuals with schizophrenia who were not on antipsychotics at the time of death (Schulmann et al., 2023), which may then have been reversed by the upregulation of *Aspa* by antipsychotics. Altogether, these findings suggest that antipsychotic treatment may enhance oligodendrocyte function or myelination in response to the frequent deficits in psychotic disorders.

### Differential Expression Related to Physiological and Cellular Stress Regulation

Chronic stress and its mechanisms have been established as a clear precursor for the onset and development of psychotic disorders, with dysregulation of the hypothalamic-pituitary-adrenal (HPA) axis serving a central role (Mondelli, 2014). The HPA axis, responsible for the regulation of the stress hormone cortisol, is commonly dysregulated in patients with psychosis, causing inappropriate cortisol secretion and impairment of the natural cortisol awakening response (Law & Clow, 2020; Stalder et al., 2025). HPA dysregulation in psychosis is theorized to contribute to functional and structural irregularities in the brain (Xenaki et al., 2024), as well as disturbed neurogenesis and cognitive function, especially affecting the PFC, hippocampus, and amygdala (Keller et al., 2017). Furthermore, chronic stress disrupts inflammatory processes within the immune system as well as oxidative stress, which may increase risk of neural disorder and psychotic illness (Rambaud et al., 2022). Antipsychotic treatment has been shown to mitigate disrupted cortisol levels within the body, normalizing HPA axis behavior, although the cortisol awakening response remains blunted (Mondelli, 2014). Antipsychotic medications can also mitigate the adverse effects of stress, including alterations in inflammatory response and dopamine signaling (Xenaki et al., 2024).

The results of our planned meta-analysis showed that six of fourteen downregulated gene pathways within the frontal cortex were associated with fear conditioning and/or stress responses, as well as two upregulated pathways. Within the non-directional results, twelve gene pathways that we identified as enriched with antipsychotic effects were related to fear conditioning and/or stress responses. This pattern was confirmed in our larger, exploratory rodent-macaque meta-analysis. The DEGs that were leading genes within these stress-related pathways included *Tdo2,* due to serotonin’s well-studied role in stress regulation (Chaouloff et al., 1999), as well as multiple genes that are likely to promote homeostasis by reducing metabolic and oxidative stress, including *Oxidative Stress Induced Growth Inhibitor 1* (*Osgin1*), *Growth Hormone* (*Gh*), and *Insulin Like Growth Factor Binding Protein 5* (*Igfbp5*).

These findings linking antipsychotics to the regulation of stress tie into a larger body of literature. Growth hormone (GH) stimulates the release and expression of insulin-like growth factor-1 (IGF-1), and *Igfbp5,* which was upregulated by antipsychotic treatment in our meta-analysis, regulates IGF signaling activity. Both GH and IGF-1 play a known role in the regulation of metabolic and oxidative stress as well as brain functions implicated in schizophrenia, such as neural development, memory formation, and dopaminergic regulation (Aguiar-Oliveira et al., 2026; Ayadi et al., 2016; Tavares et al., 2023). Both GH and IGF-1 have been linked to psychosis: mutations in *Gh* have been associated with altered prepulse inhibition in mice (Baldarelli et al., 2024; Wilson et al., 2026), and elevated IGF-1 levels have been observed in first-episode schizophrenia patients, correlating to symptom severity (Aguiar-Oliveira et al., 2026). In parallel, *Osgin1* functions as an oxidative stress-responsive gene that regulates cell apoptosis and cell proliferation, suggesting involvement in cellular stress regulation (Hussey et al., 2025).

Together, these findings may suggest that antipsychotic-associated transcriptional changes converge on pathways related to physiological and cellular stress within the brain, and reinforce the role of antipsychotics in addressing the neural effects of stress within the frontal cortex that contribute to psychosis.

### Differential Expression Related to Vasculature and the Cardiovascular System

Psychotic illnesses and antipsychotics are known to affect neurovasculature, including the blood-brain barrier (BBB), brain endothelium, and the cardiovascular system. The vascular hypothesis of schizophrenia suggests that BBB and brain endothelium dysfunction, in conjunction with environmental stressors such as hypoxia and inflammation, may contribute to schizophrenia pathophysiology (Puvogel et al., 2022). When disrupted, the BBB, a controller of central nervous system homeostasis through solute transport regulation, can drive neuroinflammation, tight junction abnormalities, and endothelial dysfunction (Puvogel et al., 2022; F. Zhang et al., 2025). Abnormalities within the BBB, including reduced capillary density and modified molecular marker expression, correlate with schizophrenia symptomology (F. Zhang et al., 2025).

With respect to antipsychotics, barrier function may also be disrupted by antipsychotic usage, as BBB endothelial cells may suffer cytotoxicity and apoptosis (F. Zhang et al., 2025). Antipsychotics can also cause metabolic, cardiac and vascular side-effects (Edinoff et al., 2022; Howell et al., 2019; Leung et al., 2012), sometimes demanding regular cardiovascular monitoring (Howell et al., 2019). These side effects enhance the risk for cardiovascular disease, which is the leading cause of death in individuals with severe mental disorders (Howell et al., 2019; Leung et al., 2012).

Our meta-analysis results converged with previous knowledge on the role of psychosis and antipsychotics on vasculature, the circulatory system, and the BBB. Twenty gene sets related to the cardiovascular system and component cell types (endothelial cells, pericytes) were enriched with differential expression within the planned meta-analysis. This pattern was confirmed in the larger, exploratory meta-analysis, which further revealed that these pathways were upregulated. The leading genes in these pathways included the upregulation of two solute carriers, *Solute Carrier Family 1 Member 5* (*Slc1a5*), which encodes an amino acid transporter Alanine-Serine-Cysteine Transporter 2 (ASCT2), and *Solute Carrier Family 22 Member 5 (Slc22a5),* which encodes a high-affinity carnitine transporter Novel Organic Cation Transporter 2 (OCTN2), accompanied by a decrease in *Fatty acid-binding protein 4* (*Fabp4*).

These findings may suggest a manner by which antipsychotics are moderating vascular changes related to psychosis. *Slc22a5* and its protein product OCTN2 are intriguing candidates because carnitine metabolite levels have been associated with both schizophrenia and cognitive improvements following antipsychotic treatment (Zhao et al., 2023) and L-carnitine is associated with cardiovascular health (Elantary & Othman, 2024). Carnitine, through its formation into acylcarnitine intermediates, plays a critical role in fatty acid β-oxidation by facilitating the transport of long-chain fatty acids into mitochondria for ATP production (Elantary & Othman, 2024). The altered acylcarnitine metabolism observed following antipsychotic treatment may reflect broader changes in mitochondrial fatty acid β-oxidation and oxidative metabolism, processes that have been linked to schizophrenia (Fizíková et al., 2023; Ni & Chung, 2020; Zhao et al., 2023). L-carnitine has further been implicated in the reduction of oxidative stress, maintenance of endothelial function, and regulation of metabolic homeostasis, overall marking it as a key player in cardiovascular health (Elantary & Othman, 2024).

Many of our top DEGs were also associated with fibroblasts, which are found perivascularly in the brain. Of these, *Sorbs2* is a compelling candidate, as mutations in *Sorbs2* produce decreased prepulse inhibition in mice (Q. Zhang et al., 2016), and caused abnormal cholesterol and cardiac function (Baldarelli et al., 2024; Wilson et al., 2026), providing another plausible link between antipsychotic therapeutic function and cardiovascular health. The upregulation of *F10* following antipsychotic treatment is also intriguing. *F10* encodes *Coagulation Factor X*, a critical protein in the blood clotting cascade. Altered plasma levels of coagulation pathway proteins have been associated with the onset of psychosis (Susai et al., 2023). Elevated plasma levels of *Coagulation Factor X* during their first episode of psychosis was previously found to predict positive antipsychotic treatment response (Susai et al., 2023).

### Strengths and Limitations

Our planned rodent meta-analysis utilized a collection of five datasets (collective *n*=68 subjects) that were identified using systematic criteria, achieving modest statistical power to determine stable estimates of gene expression regulation as a response to antipsychotic treatment. In comparison to the original studies, our meta-analysis has greater power to identify differentially expressed genes that are generalizable across antipsychotic drug class, dosage, and rodent species. Indeed, an exploratory meta-regression that included antipsychotic class as a co-variate indicated that our findings were shared across drug types.

However, our planned meta-analysis had several notable limitations that complicated the interpretation of its findings. First, the sample failed to cover a full set of characteristics that may contribute to antipsychotic treatment efficacy. Of the selected five datasets, four utilized male rodent models (details not provided for GSE4031.1), which limits any generalizability of our findings to female subjects. Research shows that sex differences have a variety of influences on the body’s responses to antipsychotics, in a manner that may also depend on hormonal balance (Seeman, 2020). Males often require a higher dosage of antipsychotic medication than females (Hoekstra et al., 2021; Seeman, 2020), and side effects are often sex-dependent as well (Hoekstra et al., 2021). Our planned meta-analysis could not address these sex differences. This limitation was particularly problematic for interpreting findings showing that the *Androgen receptor* (*Ar*), which is the primary receptor for testosterone and dihydrotestosterone, was consistently down-regulated in response to antipsychotic treatment.

Another limitation to our planned meta-analysis was that the rodent models typically received short-term treatment, with treatment durations lasting from 21 to 28 days. Past studies have shown that antipsychotic treatment in rodents shows early-onset effects, with continuous treatment progressively increasing results; effects are reversible and dependent on the duration of treatment (Li et al., 2007). Chronic antipsychotic medication usage has also been associated with brain volume change, although results are contradictory and difficult to discern from illness progression (Emsley, 2023; Lawrie, 2018). Species-specific metabolism and pharmacology may also affect the transcriptional responses to antipsychotic medications, as well as notable anatomical differences between rodent and human prefrontal cortical regions, such as the absence of granular PFC development in rodents (Preuss & Wise, 2022).

To address these limitations and increase our collective sample size to *n*=101, we ran an exploratory meta-analysis that included both the original rodent studies and data from rhesus macaques. The rhesus macaques represented both males and females, and received antipsychotic treatment for a longer duration (6 months). As non-human primates, the rhesus macaques have prefrontal anatomy that more strongly resembles humans (Preuss & Wise, 2022). Therefore, findings that were confirmed within the exploratory meta-analysis supports their generalizability, although it should be noted that the macaques still contributed only one third of the final sample size and represented results derived from a single study. Therefore, the generalizability of our findings remains an open question that should be pursued, as well as further validation.

Finally, although our meta-analysis was sufficiently powered to detect the converging effects of different drug treatments on the frontal cortex, we lacked the power to detect the differential effects of the specific drug treatments, including subgroup analyses of drug, dose, or duration-specific gene expression. We attempted to perform an exploratory meta-analysis including antipsychotic type (typical vs. atypical) as a co-variate, but it only revealed a single gene with differential expression by drug type (*Neuromedin U Receptor 1* (*Nmur1*)). Beyond broad antipsychotic drug categories, it would be especially useful to understand the divergent effects of Clozapine, in particular, on frontal cortical gene expression, as it is the only approved treatment for treatment-resistant schizophrenia (TRS) due to its strong efficacy and frequent use as a clinical indicator of treatment resistance (de Bartolomeis et al., 2022; Ying et al., 2023). Future better-powered studies should tackle these topics, as well as the tailoring of drug treatments by subject characteristics. Finally, it would be useful to explore differential effects across anatomical regions, including those likely to be mediating metabolic side effects, and to use RNA-sequencing to better characterize effects of antipsychotics on the expression of non-protein coding genes, as our meta-analysis depended heavily on datasets using older microarray platforms.

### Clinical Relevance

One notable weakness in our meta-analysis is that the included datasets solely consisted of control animals rather than models of psychosis, which means that our findings may not fully pertain to individuals in states of psychosis. However, it is worth calling out that animal models of psychosis may not necessarily serve as ideal references for antipsychotic efficacy in humans. Researchers have looked to replicate the etiologies, brain pathologies, and behavioral abnormalities behind psychotic disorders in rodent models (Young et al., 2010) by using developmental, drug-induced, lesion and genetic manipulations (Ayyar & Ravinder, 2023; Winship et al., 2019). However, it is unclear whether animals can truly model the distinctive human characteristics and experiences of psychosis, such as paranoid delusions and auditory hallucinations (Winship et al., 2019). Animal models also often fail to replicate negative symptoms of psychosis—such as alogia, blunted affect, asociality, anhedonia, and avolition (Stahl & Buckley, 2007)—impeding the ability to quantify the effect of antipsychotics on these symptoms (C. Jones et al., 2011; Winship et al., 2019). This is important because the negative symptoms of psychosis are often the most resistant to antipsychotic treatment (C. Jones et al., 2011).

Therefore, to explore the clinical relevance of our findings we compared our meta-analysis results to previous gene expression findings from post-mortem studies in human subjects with schizophrenia (Gandal et al., 2018; Schulmann et al., 2023). Although exploratory, this comparison supports the assertion that some differential expression associated with schizophrenia may represent long-term effects of antipsychotic treatment that is not easily disambiguated by post-mortem toxicology. However, the comparison overall was noisy, and suggests that both our animal meta-analysis findings and human post-mortem findings - particularly those with accompanying toxicology - may still be underpowered. Our findings may still serve as groundwork for future investigations in humans, such as comparing human postmortem brains from subjects with longer or shorter durations of antipsychotic treatment, as documented by clinical records and family interviews.

## Conclusion

In conclusion, our meta-analysis provides significant insight into the effects of antipsychotic medications on the frontal cortex. We observed a consistent molecular signature of antipsychotic treatment on the frontal cortex despite differences in the chosen antipsychotic medications, experimental procedures, overall study designs, and species. The identification of these genes helps highlight critical functional pathways that respond to antipsychotic treatment, and illuminate effects following antipsychotic administration that extend beyond classical dopaminergic signaling pathways, including genes and pathways related to serotonergic and cholinergic signalling, oligodendrocyte development and myelination, physiological and cellular stress responses, and cardiovascular function. Psychotic illnesses continue to present substantial challenges in the clinical world, and a full comprehension of the effects of antipsychotic medications on the brain may help assist medical experts and researchers achieve more targeted and effective treatment.

## CRediT statement

MRB: Conceptualization, Methodology, Software, Formal analysis, Investigation, Data Curation, Writing - Original Draft, Writing - Review & Editing, Visualization

MHH: Conceptualization, Methodology, Software, Formal Analysis, Writing - Review & Editing, Visualization, Validation, Funding Acquisition, Supervision, Project administration

EMG: Conceptualization, Methodology, Writing - Review & Editing

EIF: Conceptualization, Methodology, Writing - Review & Editing

SJW: Writing - Review & Editing, Funding acquisition

HA: Writing - Review & Editing, Supervision, Funding acquisition

## Acknowledgements

This work was completed as part of the *Brain Data Alchemy Project* and supported by the Hope for Depression Research Foundation (HDRF: HA), National Institute on Drug Abuse (NIDA U01 DA043098: HA & SJW), and the Pritzker Neuropsychiatric Disorders Research Foundation (HA & SJW). Funders and sponsors had no active role in the review.

We would like to thank Marquis Vawter for guidance regarding search terms and insight into comparisons with the crab-eating macaque data. We would like to thank Anton Schulmann for guidance and insight regarding comparisons with the findings from the rhesus macaques and human subjects with or without antipsychotics on board at the time of death. We would also like to thank Dave Krolewski and other members of the 2024 Brain Data Alchemy cohort (Sophie Mensch, Sophia Espinoza) for their insights.

## References

Aguiar-Oliveira, M. H., Boguszewski, M. C. S., Rovaris, D. L., & Donato, J. (2026). Growth Hormone and IGF-1 Actions in the Brain and Neuropsychiatric Diseases. Physiology, 41(1), 3–15. 10.1152/physiol.00009.2025

Arciniegas, D. B. (2015). Psychosis. Continuum : Lifelong Learning in Neurology, 21(3 Behavioral Neurology and Neuropsychiatry), 715–736. 10.1212/01.CON.0000466662.89908.e7

Ayadi, A. E., Zigmond, M. J., & Smith, A. D. (2016). IGF-1 protects dopamine neurons against oxidative stress: Association with changes in phosphokinases. Experimental Brain Research, 234(7), 1863–1873. 10.1007/s00221-016-4572-1

Ayyar, P., & Ravinder, J. R. (2023). Animal models for the evaluation of antipsychotic agents. Fundamental & Clinical Pharmacology, 37(3), 447–460. 10.1111/fcp.12855

Baldarelli, R. M., Smith, C. L., Ringwald, M., Richardson, J. E., Bult, C. J., & Mouse Genome Informatics Group. (2024). Mouse Genome Informatics: An integrated knowledgebase system for the laboratory mouse. Genetics, 227(1), iyae031. 10.1093/genetics/iyae031

Barrett, T., Wilhite, S. E., Ledoux, P., Evangelista, C., Kim, I. F., Tomashevsky, M., Marshall, K. A., Phillippy, K. H., Sherman, P. M., Holko, M., Yefanov, A., Lee, H., Zhang, N., Robertson, C. L., Serova, N., Davis, S., & Soboleva, A. (2013). NCBI GEO: Archive for functional genomics data sets—update. Nucleic Acids Research, 41(D1), D991–D995. 10.1093/nar/gks1193

Bilecki, W., & Maćkowiak, M. (2023). Gene Expression and Epigenetic Regulation in the Prefrontal Cortex of Schizophrenia. Genes, 14(2), 243. 10.3390/genes14020243

Bowling, H., & Santini, E. (2016). Unlocking the molecular mechanisms of antipsychotics—A new frontier for discovery. Swiss Medical Weekly, 146(27–28), Article 27–28. 10.4414/smw.2016.14314

Briles, J. J., Rosenberg, D. R., Brooks, B. A., Roberts, M. W., & Diwadkar, V. A. (2012). Review of the safety of second-generation antipsychotics: Are they really “atypically” safe for youth and adults? The Primary Care Companion for CNS Disorders, 14(3), PCC.11r01298. 10.4088/PCC.11r01298

Chaouloff, F., Berton, O., & Mormède, P. (1999). Serotonin and stress. Neuropsychopharmacology: Official Publication of the American College of Neuropsychopharmacology, 21(2 Suppl), 28S–32S. 10.1016/S0893-133X(99)00008-1

Chelban, V., Patel, N., Vandrovcova, J., Zanetti, M. N., Lynch, D. S., Ryten, M., Botía, J. A., Bello, O., Tribollet, E., Efthymiou, S., Davagnanam, I., SYNAPSE Study Group, Bashiri, F. A., Wood, N. W., Rothman, J. E., Alkuraya, F. S., & Houlden, H. (2017). Mutations in NKX6-2 Cause Progressive Spastic Ataxia and Hypomyelination. American Journal of Human Genetics, 100(6), 969–977. 10.1016/j.ajhg.2017.05.009

Chopra, S., Levi, P. T., Holmes, A., Orchard, E. R., Segal, A., Francey, S. M., O’Donoghue, B., Cropley, V. L., Nelson, B., Graham, J., Baldwin, L., Yuen, H. P., Allott, K., Alvarez-Jimenez, M., Harrigan, S., Pantelis, C., Wood, S. J., McGorry, P., & Fornito, A. (2025). Brainwide Anatomical Connectivity and Prediction of Longitudinal Outcomes in Antipsychotic-Naïve First-Episode Psychosis. Biological Psychiatry, 97(2), 157–166. 10.1016/j.biopsych.2024.07.016

Christensen, M. K., Lim, C. C. W., Saha, S., Plana-Ripoll, O., Cannon, D., Presley, F., Weye, N., Momen, N. C., Whiteford, H. A., Iburg, K. M., & McGrath, J. J. (2020). The cost of mental disorders: A systematic review. Epidemiology and Psychiatric Sciences, 29, e161. 10.1017/S204579602000075X

Christian, R., Saavedra, L., Gaynes, B. N., Sheitman, B., Wines, R. C., Jonas, D. E., Viswanathan, M., Ellis, A. R., Woodell, C., & Carey, T. S. (2012). Tables of FDA-Approved Indications for First-and Second-Generation Antipsychotics. In Future Research Needs for First- and Second-Generation Antipsychotics for Children and Young Adults [Internet]. Agency for Healthcare Research and Quality (US). https://www.ncbi.nlm.nih.gov/books/NBK84656/

Commissioner, O. of the. (2024, September 27). FDA Approves Drug with New Mechanism of Action for Treatment of Schizophrenia. FDA. FDA. https://www.fda.gov/news-events/press-announcements/fda-approves-drug-new-mechanism-action-treatment-schizophrenia

Crocker, C. E., & Tibbo, P. G. (2018). Confused Connections? Targeting White Matter to Address Treatment Resistant Schizophrenia. Frontiers in Pharmacology, 9. 10.3389/fphar.2018.01172

Davis, S., & Meltzer, P. S. (2007). GEOquery: A bridge between the Gene Expression Omnibus (GEO) and BioConductor. Bioinformatics (Oxford, England), 23(14), 1846–1847. 10.1093/bioinformatics/btm254

de Bartolomeis, A., Ciccarelli ,Mariateresa, Vellucci ,Licia, Fornaro ,Michele, Iasevoli ,Felice, & and Barone, A. (2022). Update on novel antipsychotics and pharmacological strategies for treatment-resistant schizophrenia. Expert Opinion on Pharmacotherapy, 23(18), 2035–2052. 10.1080/14656566.2022.2145884

Edinoff, A. N., Ellis, E. D., Nussdorf, L. M., Hill, T. W., Cornett, E. M., Kaye, A. M., & Kaye, A. D. (2022). Antipsychotic Polypharmacy-Related Cardiovascular Morbidity and Mortality: A Comprehensive Review. Neurology International, 14(1), 294–309. 10.3390/neurolint14010024

Elantary, R., & Othman, S. (2024). Role of L-carnitine in Cardiovascular Health: Literature Review. Cureus, 16(9), e70279. 10.7759/cureus.70279

Emsley, R. (2023). Antipsychotics and structural brain changes: Could treatment adherence explain the discrepant findings? Therapeutic Advances in Psychopharmacology, 13, 20451253231195258. 10.1177/20451253231195258

Fabrazzo, M., Cipolla, S., Camerlengo, A., Perris, F., & Catapano, F. (2022). Second-Generation Antipsychotics’ Effectiveness and Tolerability: A Review of Real-World Studies in Patients with Schizophrenia and Related Disorders. Journal of Clinical Medicine, 11(15), 4530. 10.3390/jcm11154530

Fatemi, S. H., Reutiman, T. J., Folsom, T. D., Bell, C., Nos, L., Fried, P., Pearce, D. A., Singh, S., Siderovski, D. P., Willard, F. S., & Fukuda, M. (2006). Chronic Olanzapine Treatment Causes Differential Expression of Genes in Frontal Cortex of Rats as Revealed by DNA Microarray Technique. Neuropsychopharmacology, 31(9), 1888–1899. 10.1038/sj.npp.1301002

Fizíková, I., Dragašek, J., & Račay, P. (2023). Mitochondrial Dysfunction, Altered Mitochondrial Oxygen, and Energy Metabolism Associated with the Pathogenesis of Schizophrenia. International Journal of Molecular Sciences, 24(9), 7991. 10.3390/ijms24097991

Franklin, R. J. M., & Simons, M. (2022). CNS remyelination and inflammation: From basic mechanisms to therapeutic opportunities. Neuron, 110(21), 3549–3565. 10.1016/j.neuron.2022.09.023

Gandal, M. J., Haney, J. R., Parikshak, N. N., Leppa, V., Ramaswami, G., Hartl, C., Schork, A. J., Appadurai, V., Buil, A., Werge, T. M., Liu, C., White, K. P., CommonMind Consortium, PsychENCODE Consortium, iPSYCH-BROAD Working Group, Horvath, S., & Geschwind, D. H. (2018). Shared molecular neuropathology across major psychiatric disorders parallels polygenic overlap. Science, 359(6376), 693–697. 10.1126/science.aad6469

GBD Results. (n.d.). Institute for Health Metrics and Evaluation. Retrieved April 13, 2025, from https://vizhub.healthdata.org/gbd-results

Geoghegan, E. M., Hagenauer, M. H., Hernandez, E., Espinoza, S., Flandreau, E. I., Nguyen, P. T., Santiago, A. N., Bhuiyan, M. R., Mensch, S., Watson, S. J., Akil, H., & Hen, R. (2025). The Converging Effects of Different Categories of Antidepressants on the Brain: A Systematic Meta-Analysis of Public Transcriptional Profiling Data from the Hippocampus and Cortex (p. 2025.04.21.648805). bioRxiv. 10.1101/2025.04.21.648805

Girgis, R. R., Javitch, J. A., & Lieberman, J. A. (2008). Antipsychotic drug mechanisms: Links between therapeutic effects, metabolic side effects and the insulin signaling pathway. Molecular Psychiatry, 13(10), 918–929. 10.1038/mp.2008.40

Gouvêa-Junqueira, D., Falvella, A. C. B., Antunes, A. S. L. M., Seabra, G., Brandão-Teles, C., Martins-de-Souza, D., & Crunfli, F. (2020). Novel Treatment Strategies Targeting Myelin and Oligodendrocyte Dysfunction in Schizophrenia. Frontiers in Psychiatry, 11. 10.3389/fpsyt.2020.00379

Grinchii, D., & Dremencov, E. (2020). Mechanism of Action of Atypical Antipsychotic Drugs in Mood Disorders. International Journal of Molecular Sciences, 21(24), 9532. 10.3390/ijms21249532

Hagenauer, M. H., Sannah, Y., Hebda-Bauer, E. K., Rhoads, C., O’Connor, A. M., Flandreau, E., Watson, S. J., & Akil, H. (2024). Resource: A curated database of brain-related functional gene sets (Brain.GMT). MethodsX, 13, 102788. 10.1016/j.mex.2024.102788

Hagenauer, M., Nguyen, D. M., Flandreau, E., Geoghegan, E., Mensch, S., Bhuiyan, M. R., Chennupati, L. T., Espinoza, S., Lewis, A., Drozman, A., Hughes, B. W., M.S, & Ph.D. (2024). Brain Data Alchemy Project: Meta-Analysis of Re-Analyzed Public Transcriptional Profiling Data in the Gemma Database (v.2024). https://protocols.io/view/brain-data-alchemy-project-meta-analysis-of-re-ana-dkxk4xkw

Hoekstra, S., Bartz-Johannessen, C., Sinkeviciute, I., Reitan, S. K., Kroken, R. A., Løberg, E.-M., Larsen, T. K., Rettenbacher, M., Johnsen, E., & Sommer, I. E. (2021). Sex differences in antipsychotic efficacy and side effects in schizophrenia spectrum disorder: Results from the BeSt InTro study. NPJ Schizophrenia, 7(1), 39. 10.1038/s41537-021-00170-3

Howell, S., Yarovova, E., Khwanda, A., & Rosen, S. D. (2019). Cardiovascular effects of psychotic illnesses and antipsychotic therapy. Heart (British Cardiac Society), 105(24), 1852–1859. 10.1136/heartjnl-2017-312107

Huhn, M., Nikolakopoulou, A., Schneider-Thoma, J., Krause, M., Samara, M., Peter, N., Arndt, T., Bäckers, L., Rothe, P., Cipriani, A., Davis, J., Salanti, G., & Leucht, S. (2019). Comparative efficacy and tolerability of 32 oral antipsychotics for the acute treatment of adults with multi-episode schizophrenia: A systematic review and network meta-analysis. The Lancet, 394(10202), 939–951. 10.1016/S0140-6736(19)31135-3

Hussey, G., Royster, M., Vaidy, N., Culkin, M., & Saha, M. S. (2025). The Osgin Gene Family: Underexplored Yet Essential Mediators of Oxidative Stress. Biomolecules, 15(3), 409. 10.3390/biom15030409

Ibi, D., de la Fuente Revenga, M., Kezunovic, N., Muguruza, C., Saunders, J. M., Gaitonde, S. A., Moreno, J. L., Ijaz, M. K., Santosh, V., Kozlenkov, A., Holloway, T., Seto, J., García-Bea, A., Kurita, M., Mosley, G. E., Jiang, Y., Christoffel, D. J., Callado, L. F., Russo, S. J., … González-Maeso, J. (2017). Antipsychotic-induced Hdac2 transcription via NF-κB leads to synaptic and cognitive side effects. Nature Neuroscience, 20(9), 1247–1259. 10.1038/nn.4616

Jones, C., Watson, D., & Fone, K. (2011). Animal models of schizophrenia. British Journal of Pharmacology, 164(4), 1162–1194. 10.1111/j.1476-5381.2011.01386.x

Jones, D. T., & Graff-Radford, J. (2021). Executive Dysfunction and the Prefrontal Cortex. CONTINUUM: Lifelong Learning in Neurology, 27(6), 1586–1601. 10.1212/CON.0000000000001009

Julayanont, P., & Suryadevara, U. (2021). Psychosis. CONTINUUM: Lifelong Learning in Neurology, 27(6), 1682. 10.1212/CON.0000000000001013

Keller, J., Gomez, R., Williams, G., Lembke, A., Lazzeroni, L., Murphy, G. M., & Schatzberg, A. F. (2017). HPA Axis in Major Depression: Cortisol, Clinical Symptomatology, and Genetic Variation Predict Cognition. Molecular Psychiatry, 22(4), 527–536. 10.1038/mp.2016.120

Kondo, M. A., Tajinda, K., Colantuoni, C., Hiyama, H., Seshadri, S., Huang, B., Pou, S., Furukori, K., Hookway, C., Jaaro-Peled, H., Kano, S. -i, Matsuoka, N., Harada, K., Ni, K., Pevsner, J., & Sawa, A. (2013). Unique pharmacological actions of atypical neuroleptic quetiapine: Possible role in cell cycle/fate control. Translational Psychiatry, 3(4), e243–e243. 10.1038/tp.2013.19

Korotkevich, G., Sukhov, V., Budin, N., Shpak, B., Artyomov, M. N., & Sergushichev, A. (2021). Fast gene set enrichment analysis (p. 060012). bioRxiv. 10.1101/060012

Lanz, T. A., Reinhart, V., Sheehan, M. J., Rizzo, S. J. S., Bove, S. E., James, L. C., Volfson, D., Lewis, D. A., & Kleiman, R. J. (2019). Postmortem transcriptional profiling reveals widespread increase in inflammation in schizophrenia: A comparison of prefrontal cortex, striatum, and hippocampus among matched tetrads of controls with subjects diagnosed with schizophrenia, bipolar or major depressive disorder. Translational Psychiatry, 9(1), 1–13. 10.1038/s41398-019-0492-8

Law, R., & Clow, A. (2020). Stress, the cortisol awakening response and cognitive function. International Review of Neurobiology, 150, 187–217. 10.1016/bs.irn.2020.01.001

Lawrie, S. M. (2018). Are structural brain changes in schizophrenia related to antipsychotic medication? A narrative review of the evidence from a clinical perspective. Therapeutic Advances in Psychopharmacology, 8(11), 319–326. 10.1177/2045125318782306

Leucht, S., Priller, J., & Davis, J. M. (2024). Antipsychotic Drugs: A Concise Review of History, Classification, Indications, Mechanism, Efficacy, Side Effects, Dosing, and Clinical Application. American Journal of Psychiatry, 181(10), 865–878. 10.1176/appi.ajp.20240738

Leung, J. Y. T., Barr, A. M., Procyshyn, R. M., Honer, W. G., & Pang, C. C. Y. (2012). Cardiovascular side-effects of antipsychotic drugs: The role of the autonomic nervous system. Pharmacology & Therapeutics, 135(2), 113–122. 10.1016/j.pharmthera.2012.04.003

Li, M., Fletcher, P. J., & Kapur, S. (2007). Time Course of the Antipsychotic Effect and the Underlying Behavioral Mechanisms. Neuropsychopharmacology, 32(2), 263–272. 10.1038/sj.npp.1301110

Lieberman, J. A., & First, M. B. (2018). Psychotic Disorders. New England Journal of Medicine, 379(3), 270–280. 10.1056/NEJMra1801490

Lim, N., Tesar, S., Belmadani, M., Poirier-Morency, G., Mancarci, B. O., Sicherman, J., Jacobson, M., Leong, J., Tan, P., & Pavlidis, P. (2021). Curation of over 10 000 transcriptomic studies to enable data reuse. Database, 2021, baab006. 10.1093/database/baab006

Martin, M. V., Mirnics, K., Nisenbaum, L. K., & Vawter, M. P. (2015). Olanzapine Reversed Brain Gene Expression Changes Induced by Phencyclidine Treatment in Non-Human Primates. Molecular Neuropsychiatry, 1(2), 82–93. 10.1159/000430786

Meltzer, H., & Massey, B. (2011). The role of serotonin receptors in the action of atypical antipsychotic drugs. Current Opinion in Pharmacology, Neurosciences, 11(1), 59–67. 10.1016/j.coph.2011.02.007

Meltzer, H. Y. (2013). Update on Typical and Atypical Antipsychotic Drugs. Annual Review of Medicine, 64(1), 393–406. 10.1146/annurev-med-050911-161504

Mighdoll, M. I., Tao, R., Kleinman, J. E., & Hyde, T. M. (2015). Myelin, myelin-related disorders, and psychosis. Schizophrenia Research, White Matter Pathology, 161(1), 85–93. 10.1016/j.schres.2014.09.040

Mondelli, V. (2014). From stress to psychosis: Whom, how, when and why? Epidemiology and Psychiatric Sciences, 23(3), 215–218. 10.1017/S204579601400033X

Morris, B. J. (2026). Parsing the neuroanatomy of schizophrenia to enhance the translational validity of preclinical models—A multidisciplinary perspective. Progress in Neuro-Psychopharmacology and Biological Psychiatry, 145, 111628. 10.1016/j.pnpbp.2026.111628

Mudunuri, U., Che, A., Yi, M., & Stephens, R. M. (2009). bioDBnet: The biological database network. Bioinformatics, 25(4), 555–556. 10.1093/bioinformatics/btn654

Ni, P., & Chung, S. (2020). Mitochondrial Dysfunction in Schizophrenia. BioEssays, 42(6), 1900202. 10.1002/bies.201900202

Olivas-Bernal, C. A., Vargas-Albores, F., Garibay-Valdez, E., Cicala, F., & Martínez-Porchas, M. (2026). Transcriptomic Meta-Analysis as a Framework for Robust Cross-Study Biological Inference. International Journal of Molecular Sciences, 27(11). 10.3390/ijms27114674

Osugo, M., Whitehurst, T., Erritzoe, D., Carr, R., Ashok, A. H., Maccioni, L., Onwordi, E. C., Rutigliano, G., Rahaman, N., Arumuham, A., de Marvao, A., Gunn, R. N., Rabiner, E. A., Marques, T. R., Veronese, M., & Howes, O. D. (2026). Role of Serotonin in the Neurobiology of Schizophrenia and Association With Negative Symptoms. JAMA Psychiatry, 83(2), 185–195. 10.1001/jamapsychiatry.2025.3430

Peters, B. D., & Karlsgodt, K. H. (2015). WHITE MATTER DEVELOPMENT IN THE EARLY STAGES OF PSYCHOSIS. Schizophrenia Research, 161(1), 61–69. 10.1016/j.schres.2014.05.021

Pollard, K. S., Dudoit, S., & van der Laan, M. J. (2005). Multiple Testing Procedures: The multtest Package and Applications to Genomics. In R. Gentleman, V. J. Carey, W. Huber, R. A. Irizarry, & S. Dudoit (Eds.), Bioinformatics and Computational Biology Solutions Using R and Bioconductor (pp. 251–272). Springer. 10.1007/0-387-29362-0_15

Posit Team. (2025). RStudio: Integrated development environment for R (Version 2024.12.1.563) [Computer software]. Posit Software, PBC. https://posit.co/

Preuss, T. M., & Wise, S. P. (2022). Evolution of prefrontal cortex. Neuropsychopharmacology, 47(1), 3–19. 10.1038/s41386-021-01076-5

Puvogel, S., Palma, V., & Sommer, I. E. C. (2022). Brain vasculature disturbance in schizophrenia. Current Opinion in Psychiatry, 35(3), 146. 10.1097/YCO.0000000000000789

R Core Team. (2024). R: A language and environment for statistical computing (Version 4.4.1) [Computer software]. R Foundation for Statistical Computing. https://www.R-project.org

Rambaud, V., Marzo, A., & Chaumette, B. (2022). Oxidative Stress and Emergence of Psychosis. Antioxidants, 11(10), 1870. 10.3390/antiox11101870

Rhoads, C. A., Hagenauer, M. H., Xiong, J., Hernandez, E., Nguyen, D. M., Saffron, A., Flandreau, E., Watson, S., & Akil, H. (2025). A Meta-Analysis of the Effects of Acute Sleep Deprivation on the Cortical Transcriptome in Rodent Models. Journal of Sleep Research, e70205. 10.1111/jsr.70205

Ritchie, M. E., Phipson, B., Wu, D., Hu, Y., Law, C. W., Shi, W., & Smyth, G. K. (2015). Limma powers differential expression analyses for RNA-sequencing and microarray studies. Nucleic Acids Research, 43(7), e47. 10.1093/nar/gkv007

Romeo, B., Willaime, L., Rari, E., Benyamina, A., & Martelli, C. (2023). Efficacy of 5-HT2A antagonists on negative symptoms in patients with schizophrenia: A meta-analysis. Psychiatry Research, 321, 115104. 10.1016/j.psychres.2023.115104

RxClass. (n.d.). Retrieved April 13, 2025, from https://mor.nlm.nih.gov/RxClass/search?query=N0000178316%7CTC&searchBy=class&sourceIds=&drugSources=ATC1-4%7CATCPROD%2CEPC%7CDAILYMED%2CDISEASE%7CMEDRT%2CCHEM%7CDAILYMED%2CMOA%7CDAILYMED%2CPE%7CDAILYMED%2CPK%7CMEDRT%2CVA%7CVA%2CTC%7CFMTSME%2CDISPOS%7CSNOMEDCT%2CSTRUCT%7CSNOMEDCT%2CTHERAP%7CSNOMEDCT%2CSCHEDULE%7CRXNORM

Saint-Georges, Z., MacDonald, J., Al-Khalili, R., Hamati, R., Solmi, M., Keshavan, M. S., Tuominen, L., & Guimond, S. (2025). Cholinergic system in schizophrenia: A systematic review and meta-analysis. Molecular Psychiatry, 30(7), 3301–3315. 10.1038/s41380-025-03023-y

Schulmann, A., Marenco, S., Vawter, M. P., Akula, N., Limon, A., Mandal, A., Auluck, P. K., Patel, Y., Lipska, B. K., & McMahon, F. J. (2023). Antipsychotic drug use complicates assessment of gene expression changes associated with schizophrenia. Translational Psychiatry, 13(1), 93. 10.1038/s41398-023-02392-8

Seeman, M. V. (2020). Men and women respond differently to antipsychotic drugs. Neuropharmacology, 163, 107631. 10.1016/j.neuropharm.2019.05.008

Siafis, S., Wu, H., Wang, D., Burschinski, A., Nomura, N., Takeuchi, H., Schneider-Thoma, J., Davis, J. M., & Leucht, S. (2023). Antipsychotic dose, dopamine D2 receptor occupancy and extrapyramidal side-effects: A systematic review and dose-response meta-analysis. Molecular Psychiatry, 28(8), 3267–3277. 10.1038/s41380-023-02203-y

Smyth, G. K. (2005). limma: Linear Models for Microarray Data. In R. Gentleman, V. J. Carey, W. Huber, R. A. Irizarry, & S. Dudoit (Eds.), Bioinformatics and Computational Biology Solutions Using R and Bioconductor (pp. 397–420). Springer. 10.1007/0-387-29362-0_23

Stahl, S. M. (2018). Beyond the dopamine hypothesis of schizophrenia to three neural networks of psychosis: Dopamine, serotonin, and glutamate. CNS Spectrums, 23(3), 187–191. 10.1017/S1092852918001013

Stahl, S. M., & Buckley, P. F. (2007). Negative symptoms of schizophrenia: A problem that will not go away. Acta Psychiatrica Scandinavica, 115(1), 4–11. 10.1111/j.1600-0447.2006.00947.x

Stalder, T., Oster, H., Abelson, J. L., Huthsteiner, K., Klucken, T., & Clow, A. (2025). The Cortisol Awakening Response: Regulation and Functional Significance. Endocrine Reviews, 46(1), 43–59. 10.1210/endrev/bnae024

Subramanian, A., Tamayo, P., Mootha, V. K., Mukherjee, S., Ebert, B. L., Gillette, M. A., Paulovich, A., Pomeroy, S. L., Golub, T. R., Lander, E. S., & Mesirov, J. P. (2005). Gene set enrichment analysis: A knowledge-based approach for interpreting genome-wide expression profiles. Proceedings of the National Academy of Sciences, 102(43), 15545–15550. 10.1073/pnas.0506580102

Susai, S. R., Föcking, M., Mongan, D., Heurich, M., Coutts, F., Egerton, A., Whetton, T., Winter-van Rossum, I., Unwin, R. D., Pollak, T. A., Weiser, M., Leboyer, M., Rujescu, D., Byrne, J. F., Gifford, G. W., Dazzan, P., Koutsouleris, N., Kahn, R. S., Cotter, D. R., & McGuire, P. (2023). Association of Complement and Coagulation Pathway Proteins With Treatment Response in First-Episode Psychosis: A Longitudinal Analysis of the OPTiMiSE Clinical Trial. Schizophrenia Bulletin, 49(4), 893–902. 10.1093/schbul/sbac201

Tavares, M. R., Frazao, R., & Donato, J. (2023). Understanding the role of growth hormone in situations of metabolic stress. The Journal of Endocrinology, 256(1), e220159. 10.1530/JOE-22-0159

Viechtbauer, W. (2010). Conducting Meta-Analyses in R with the metafor Package. Journal of Statistical Software, 36, 1–48. 10.18637/jss.v036.i03

Weinstein, J. J., Moeller, S. J., Perlman, G., Gil, R., Van Snellenberg, J. X., Wengler, K., Meng, J., Slifstein, M., & Abi-Dargham, A. (2024). Imaging the Vesicular Acetylcholine Transporter in Schizophrenia: A Positron Emission Tomography Study Using [18F]-VAT. Biological Psychiatry, 96(5), 352–364. 10.1016/j.biopsych.2024.01.019

Wilson, R., Bülbül Ataç, T., Cheng, T. K., Frost, A., Güneş, O., Kan, M., Keskivali-Bond, P., López Gómez, F., McLaughlin, J., Mucha, J., Munava, T., Oliveira, C., Pava, D., Peña Estrada, J. F., Selkirk, E., Vardal, B., Wells, S., Cacheiro, P., Smedley, D., & Parkinson, H. (2026). International Mouse Phenotyping Consortium Portal: Facilitating investigation of gene function and providing insights into human disease. Nucleic Acids Research, 54(D1), D1133–D1142. 10.1093/nar/gkaf1148

Winship, I. R., Dursun, S. M., Baker, G. B., Balista, P. A., Kandratavicius, L., Maia-de-Oliveira, J. P., Hallak, J., & Howland, J. G. (2019). An Overview of Animal Models Related to Schizophrenia. The Canadian Journal of Psychiatry, 64(1), 5–17. 10.1177/0706743718773728

Xenaki, L.-A., Dimitrakopoulos, S., Selakovic, M., & Stefanis, N. (2024). Stress, Environment and Early Psychosis. Current Neuropharmacology, 22(3), 437–460. 10.2174/1570159X21666230817153631

Xiong, J., Hagenauer, M. H., Rhoads, C. A., Flandreau, E., Rempel-Clower, N., Hernandez, E., Nguyen, D. M., Saffron, A., Duan, T., Watson, S., & Akil, H. (2026). A Meta-Analysis of the Effects of Chronic Stress on the Prefrontal Transcriptome in Animal Models and Convergence With Existing Human Data. Brain and Behavior, 16(1), e71197. 10.1002/brb3.71197

Ying, J., Chew, Q. H., McIntyre, R. S., & Sim, K. (2023). Treatment-Resistant Schizophrenia, Clozapine Resistance, Genetic Associations, and Implications for Precision Psychiatry: A Scoping Review. Genes, 14(3), 689. 10.3390/genes14030689

Yohn, S. E., Weiden, P. J., Felder, C. C., & Stahl, S. M. (2022). Muscarinic acetylcholine receptors for psychotic disorders: Bench-side to clinic. Trends in Pharmacological Sciences, 43(12), 1098–1112. 10.1016/j.tips.2022.09.006

Young, J. W., Zhou, X., & Geyer, M. A. (2010). Animal models of schizophrenia. Current Topics in Behavioral Neurosciences, 4, 391–433. 10.1007/7854_2010_62

Zhang, F., Zhang, J., Wang, X., Han, M., Fei, Y., & Wang, J. (2025). Blood–Brain Barrier Disruption in Schizophrenia: Insights, Mechanisms, and Future Directions. International Journal of Molecular Sciences, 26(3), 873. 10.3390/ijms26030873

Zhang, Q., Gao, X., Li, C., Feliciano, C., Wang, D., Zhou, D., Mei, Y., Monteiro, P., Anand, M., Itohara, S., Dong, X., Fu, Z., & Feng, G. (2016). Impaired Dendritic Development and Memory in Sorbs2 Knock-Out Mice. The Journal of Neuroscience: The Official Journal of the Society for Neuroscience, 36(7), 2247–2260. 10.1523/JNEUROSCI.2528-15.2016

Zhao, L., Liu, H., Wang, W., Wang, Y., Xiu, M., & Li, S. (2023). Carnitine metabolites and cognitive improvement in patients with schizophrenia treated with olanzapine: A prospective longitudinal study. Frontiers in Pharmacology, 14, 1255501. 10.3389/fphar.2023.1255501

Zhou, X., & Stephens, M. (2012). Genome-wide efficient mixed-model analysis for association studies. Nature Genetics, 44(7), 821–824. 10.1038/ng.2310 10.1038/ng.2310

