## Supplemental Tables, Figures, and Legends for "A Meta-Analysis of the Converging Effects of Different Classes of Antipsychotics on the Frontal Cortex Transcriptome in Laboratory Rodents and Non-Human Primates"

#### Supplemental Table Legends

***Table S1. The input used for the meta-analyses.*** *This .xlsx file includes 3 worksheets: 1) The worksheet "Column Definitions" contains the definitions for each of the statistical worksheets and columns. 2) The worksheet "Log2FCs" provides the average log(2) fold changes for each gene for each of the antipsychotic treatment vs. control comparisons included in either the planned rodent meta-analysis or exploratory rodent-macque meta-analysis. In order to be included within either the planned rodent meta-analysis or exploratory rodent-macaque meta-analysis, a gene needed to be present within at least 7 of the available antipsychotic treatment vs. control comparisons. The gene annotation for each of the 13,121 rows is provided for each relevant species (mouse, rat, macaque (with human)), as well as information about the evidence supporting gene orthology across species. The antipsychotic treatment vs. control comparisons are identified by a dataset identifier (either gene expression omnibus accession# or research consortium) and treatment name. The Log2FCs were averaged if there were multiple results available for any particular gene (as identified by mouse Entrez ID, rat Entrez ID, or the mouse Entrez ID for the respective macaque/human ortholog). 3) The worksheet "SamplingVariances" provides the sampling variances for each gene to accompany the Log2FCs in worksheet #1 (13,121 genes). The sampling variances were calculated as the square of the standard error (Log2FC/T-statistic). The standard errors were averaged if there were multiple results available for any particular gene (as identified by mouse Entrez ID, rat Entrez ID, or the mouse Entrez ID for the respective macaque/human ortholog).*

***Table S2. Planned meta-analysis results: The full results from the planned meta-analysis of antipsychotic effects on the frontal cortex of rodents (12,215 genes, 12,190 stable meta-analysis estimates).*** *This .xlsx file includes two worksheets: 1) The worksheet "ColumnDefinitions" provides the definitions for the variables present in each column in "PlannedRodentMetaAnalysisOutput". 1) The worksheet "PlannedRodentMetaAnalysisOutput" provides the full meta-analysis results, with each row representing the results for one gene, and each column providing either gene annotation or meta-analysis statistical output. The results are ordered by p-value, so that the top rows in the worksheet are the genes with the smallest p-values.*

***Table S3. Planned meta-analysis results: All 45 genes consistently upregulated in the frontal cortex following antipsychotic treatment in rodents (FDR<0.05).*** *Results are ordered by estimated Log2FC (Log2 Fold Change of Antipsychotic treatment vs. Control); a higher Log2FC indicates upregulation of the gene within the frontal cortex following antipsychotic treatment compared to the control group. Given gene symbols are the mouse symbol for the respective gene. Definitions: Log2FC = Log2 Fold Change, SE = Standard Error, CI_lb=confidence interval lower bound, CI_ub=confidence interval upper bound, p-value = nominal p-value, FDR = False Discovery Rate (q-value). The #Comparisons represents the number of treatments that included measurements for the gene following quality control. The "Confirmed?" column indicates whether a gene continued to show differential expression in the larger exploratory meta-analysis drawing from data from both rodents and macaques - a single asterisk (*) indicates that the gene continued to show significant differential expression (FDR<0.05) in the larger exploratory meta-analysis, regardless of whether macaque data was available, whereas double asterisks (**) indicate that the gene was both significant and had rhesus macaque data available. Full results for all genes can be found in* ***Table S2****.*

| Gene Symbol | Mouse Entrez Gene ID | Log2FC estimate | SE | pval | CI_lb | CI_ub | #Comparisons | FDR | Confirmed? |
|---|---|---|---|---|---|---|---|---|---|
| ***Nkx6-2*** | 14912 | 0.288 | 0.0608 | 2.06E-06 | 0.169 | 0.408 | 8 | 5.03E-03 | ** |
| ***Plekhh1*** | 211945 | 0.274 | 0.0697 | 8.40E-05 | 0.137 | 0.410 | 8 | 2.44E-02 | |
| ***Gjb1*** | 14618 | 0.271 | 0.0588 | 4.04E-06 | 0.156 | 0.386 | 8 | 7.04E-03 | * |
| ***Mfap2*** | 17150 | 0.246 | 0.0390 | 2.99E-10 | 0.169 | 0.322 | 8 | 1.83E-06 | ** |
| ***Igfbp5*** | 16011 | 0.238 | 0.0564 | 2.47E-05 | 0.127 | 0.348 | 8 | 1.40E-02 | ** |
| ***Mdc1*** | 240087 | 0.228 | 0.0508 | 7.59E-06 | 0.128 | 0.327 | 8 | 7.27E-03 | |
| ***Cdc42ep2*** | 104252 | 0.210 | 0.0470 | 8.33E-06 | 0.117 | 0.302 | 8 | 7.27E-03 | |
| ***Prok1*** | 246691 | 0.205 | 0.0519 | 7.66E-05 | 0.104 | 0.307 | 7 | 2.34E-02 | * |
| ***Ccdc71*** | 72454 | 0.200 | 0.0384 | 1.87E-07 | 0.125 | 0.276 | 7 | 5.70E-04 | ** |
| ***Gh*** | 14599 | 0.192 | 0.0496 | 1.11E-04 | 0.095 | 0.289 | 8 | 2.87E-02 | * |
| ***Osgin1*** | 71839 | 0.186 | 0.0492 | 1.59E-04 | 0.089 | 0.282 | 8 | 3.60E-02 | ** |
| ***Pola2*** | 18969 | 0.186 | 0.0408 | 5.49E-06 | 0.106 | 0.266 | 8 | 7.27E-03 | |
| ***Cpne7*** | 102278 | 0.185 | 0.0436 | 2.25E-05 | 0.099 | 0.271 | 8 | 1.40E-02 | |
| ***Rlbp1*** | 19771 | 0.183 | 0.0411 | 8.13E-06 | 0.103 | 0.264 | 8 | 7.27E-03 | |
| ***Aspa*** | 11484 | 0.182 | 0.0432 | 2.41E-05 | 0.098 | 0.267 | 8 | 1.40E-02 | ** |
| ***Slc18a3*** | 20508 | 0.182 | 0.0457 | 6.85E-05 | 0.092 | 0.272 | 8 | 2.34E-02 | * |
| ***Tdo2*** | 56720 | 0.181 | 0.0404 | 7.78E-06 | 0.102 | 0.260 | 8 | 7.27E-03 | * |
| ***1700010I14Rik*** | 66931 | 0.172 | 0.0412 | 3.10E-05 | 0.091 | 0.253 | 8 | 1.40E-02 | * |
| ***Gpr84*** | 80910 | 0.171 | 0.0417 | 4.27E-05 | 0.089 | 0.253 | 8 | 1.80E-02 | * |
| ***Slc14a1*** | 108052 | 0.170 | 0.0376 | 6.27E-06 | 0.096 | 0.243 | 8 | 7.27E-03 | ** |
| ***Slc1a5*** | 20514 | 0.159 | 0.0371 | 1.95E-05 | 0.086 | 0.231 | 8 | 1.40E-02 | * |
| ***Tppp3*** | 67971 | 0.159 | 0.0400 | 7.28E-05 | 0.080 | 0.237 | 8 | 2.34E-02 | |
| ***Tspo*** | 12257 | 0.159 | 0.0378 | 2.70E-05 | 0.085 | 0.233 | 8 | 1.40E-02 | |
| ***Slc9a8*** | 77031 | 0.155 | 0.0392 | 7.14E-05 | 0.079 | 0.232 | 7 | 2.34E-02 | |
| ***Plekhh3*** | 217198 | 0.148 | 0.0386 | 1.23E-04 | 0.073 | 0.224 | 8 | 3.01E-02 | |
| ***Sgcg*** | 24053 | 0.148 | 0.0384 | 1.23E-04 | 0.072 | 0.223 | 7 | 3.01E-02 | |
| ***F10*** | 14058 | 0.145 | 0.0382 | 1.45E-04 | 0.070 | 0.220 | 7 | 3.47E-02 | ** |
| ***Cers2*** | 76893 | 0.145 | 0.0362 | 6.42E-05 | 0.074 | 0.215 | 8 | 2.31E-02 | |
| ***Cmtm3*** | 68119 | 0.144 | 0.0360 | 5.89E-05 | 0.074 | 0.215 | 8 | 2.25E-02 | ** |
| ***Mtmr14*** | 97287 | 0.142 | 0.0302 | 2.78E-06 | 0.082 | 0.201 | 8 | 5.67E-03 | |
| ***Gtse1*** | 29870 | 0.142 | 0.0366 | 1.08E-04 | 0.070 | 0.213 | 8 | 2.87E-02 | * |
| ***Slc22a5*** | 20520 | 0.141 | 0.0337 | 3.04E-05 | 0.075 | 0.207 | 8 | 1.40E-02 | ** |
| ***Mrgprf*** | 211577 | 0.140 | 0.0353 | 6.93E-05 | 0.071 | 0.209 | 8 | 2.34E-02 | ** |
| ***Plekhj1*** | 78670 | 0.139 | 0.0355 | 8.91E-05 | 0.070 | 0.209 | 8 | 2.47E-02 | |
| ***Actl6b*** | 83766 | 0.127 | 0.0302 | 2.62E-05 | 0.068 | 0.186 | 8 | 1.40E-02 | |
| ***Rbp7*** | 63954 | 0.123 | 0.0329 | 1.82E-04 | 0.059 | 0.187 | 7 | 3.96E-02 | * |
| ***Tmem134*** | 66990 | 0.118 | 0.0278 | 2.38E-05 | 0.063 | 0.172 | 8 | 1.40E-02 | ** |
| ***Tbc1d17*** | 233204 | 0.117 | 0.0264 | 9.77E-06 | 0.065 | 0.168 | 8 | 7.96E-03 | |
| ***Fam234a*** | 106581 | 0.116 | 0.0306 | 1.56E-04 | 0.056 | 0.176 | 8 | 3.60E-02 | * |
| ***Cd180*** | 17079 | 0.110 | 0.0280 | 8.36E-05 | 0.055 | 0.165 | 8 | 2.44E-02 | ** |
| ***Hnrnpl*** | 15388 | 0.109 | 0.0291 | 1.69E-04 | 0.052 | 0.166 | 8 | 3.76E-02 | * |
| ***Pcyt2*** | 68671 | 0.100 | 0.0269 | 1.85E-04 | 0.048 | 0.153 | 8 | 3.96E-02 | |
| ***Kctd2*** | 70382 | 0.100 | 0.0237 | 2.42E-05 | 0.054 | 0.146 | 7 | 1.40E-02 | |
| ***Polr2c*** | 20021 | 0.100 | 0.0144 | 4.50E-12 | 0.071 | 0.128 | 8 | 5.49E-08 | |
| ***Txndc15*** | 69672 | 0.069 | 0.0182 | 1.56E-04 | 0.033 | 0.104 | 8 | 3.60E-02 | |

***Table S4. Planned meta-analysis results: All 18 genes consistently downregulated in the frontal cortex following antipsychotic treatment in rodents (FDR<0.05).*** *Table follows the conventions of* ***Table S3****. Results are ordered by estimated Log2FC (Log2 Fold Change of Antipsychotic treatment vs. Control); a lower Log2FC indicates downregulation of the gene within the frontal cortex following antipsychotic treatment compared to the control group. Full results for all genes can be found in* ***Table S2****.*

| Gene Symbol | Mouse Entrez Gene ID | Log2FC estimate | SE | pval | CI_lb | CI_ub | #Comparisons | FDR | Confirmed? |
|---|---|---|---|---|---|---|---|---|---|
| ***Ms4a8a*** | 64381 | -0.248 | 0.0427 | 6.28E-09 | -0.331 | -0.164 | 7 | 2.56E-05 | * |
| ***Plcg2*** | 234779 | -0.230 | 0.0573 | 5.86E-05 | -0.343 | -0.118 | 8 | 2.25E-02 | |
| ***Sowaha*** | 237761 | -0.203 | 0.0484 | 2.89E-05 | -0.298 | -0.108 | 7 | 1.40E-02 | |
| ***Atp8a2*** | 50769 | -0.192 | 0.0499 | 1.20E-04 | -0.290 | -0.094 | 7 | 3.01E-02 | |
| ***Gpr45*** | 93690 | -0.179 | 0.0482 | 2.08E-04 | -0.273 | -0.084 | 8 | 4.17E-02 | |
| ***Fabp4*** | 11770 | -0.168 | 0.0417 | 5.36E-05 | -0.250 | -0.087 | 7 | 2.18E-02 | ** |
| ***Ica1l*** | 70375 | -0.160 | 0.0435 | 2.34E-04 | -0.245 | -0.075 | 7 | 4.60E-02 | |
| ***Tnfsf10*** | 22035 | -0.148 | 0.0403 | 2.47E-04 | -0.227 | -0.069 | 7 | 4.79E-02 | |
| ***Ar*** | 11835 | -0.137 | 0.0369 | 2.02E-04 | -0.210 | -0.065 | 8 | 4.17E-02 | ** |
| ***Tasp1*** | 75812 | -0.136 | 0.0347 | 8.65E-05 | -0.204 | -0.068 | 8 | 2.46E-02 | ** |
| ***Nt5c*** | 50773 | -0.136 | 0.0349 | 1.01E-04 | -0.204 | -0.067 | 8 | 2.73E-02 | |
| ***Nol4l*** | 329540 | -0.123 | 0.0307 | 6.29E-05 | -0.183 | -0.063 | 7 | 2.31E-02 | |
| ***Ncr1*** | 17086 | -0.120 | 0.0290 | 3.44E-05 | -0.177 | -0.063 | 7 | 1.50E-02 | * |
| ***Sorbs2*** | 234214 | -0.118 | 0.0260 | 5.94E-06 | -0.169 | -0.067 | 8 | 7.27E-03 | |
| ***Nudt7*** | 67528 | -0.117 | 0.0315 | 2.06E-04 | -0.179 | -0.055 | 8 | 4.17E-02 | ** |
| ***Cdk5r1*** | 12569 | -0.104 | 0.0280 | 2.08E-04 | -0.159 | -0.049 | 8 | 4.17E-02 | |
| ***B4galnt1*** | 14421 | -0.104 | 0.0262 | 7.64E-05 | -0.155 | -0.052 | 8 | 2.34E-02 | |
| ***Pprc1*** | 226169 | -0.089 | 0.0213 | 2.92E-05 | -0.131 | -0.047 | 8 | 1.40E-02 | |

***Table S5. Exploratory results from a meta-regression including antipsychotic type (second-generation vs. first-generation) as a co-variate (12,215 genes, 12,203 stable meta-analysis estimates).*** *This .xlsx file includes two worksheets: 1) The worksheet "ColumnDefinitions" provides the definitions for the variables present in each column in "ExploratoryMetaRegressionResults". 1) The worksheet "ExploratoryMetaRegressionResults" provides the full meta-analysis results, with each row representing the results for one gene, and each column providing either gene annotation or meta-analysis statistical output. The results are ordered by p-value for the overall effect of the covariate antipsychotic type, so that the top rows in the worksheet are the genes with the smallest p-values.*

***Table S6. 39 genes showed significant evidence of publication bias when considering the full meta-analysis output (Egger's test FDR<0.05).*** *The Egger's regression test is used to detect funnel plot asymmetry, which is a commonly used indicator of publication bias. Rows in the table are ordered by the p-value from the Egger's regression test ("Egger Pval"), provided with the associated Z statistic ("Egger Zstat") and false discovery rate ("Egger FDR"). Other columns follow the conventions used in* ***Table S3*** *and* ***Table S4****, but with "AP" listed before each statistic to signify "Antipsychotic Effect". Full results for all genes can be found in* ***Table S2****.*

| Gene Symbol | Mouse Entrez Gene ID | AP Log2FC estimate | AP SE | AP pval | AP CI_lb | AP CI_ub | # Comparisons | AP FDR | Egger Zstat | Egger Pval | Egger FDR |
|---|---|---|---|---|---|---|---|---|---|---|---|
| ***Tepsin*** | *78777* | *0.28* | *0.32* | *3.69E-01* | *-0.34* | *0.90* | *8* | *8.26E-01* | *5.30* | *1.15E-07* | *1.24E-03* |
| ***Strip2*** | *320609* | *-0.04* | *0.12* | *7.39E-01* | *-0.28* | *0.20* | *7* | *9.51E-01* | *-5.13* | *2.87E-07* | *1.24E-03* |
| ***Utp14a*** | *72554* | *-0.08* | *0.06* | *1.93E-01* | *-0.20* | *0.04* | *8* | *7.01E-01* | *-5.12* | *3.04E-07* | *1.24E-03* |
| ***Tbx18*** | *76365* | *0.00* | *0.12* | *9.91E-01* | *-0.24* | *0.24* | *8* | *1.00E+00* | *4.96* | *6.94E-07* | *2.12E-03* |
| ***Polm*** | *54125* | *0.02* | *0.07* | *7.13E-01* | *-0.11* | *0.15* | *8* | *9.48E-01* | *4.75* | *1.99E-06* | *4.86E-03* |
| ***Tradd*** | *71609* | *-0.26* | *0.14* | *6.54E-02* | *-0.54* | *0.02* | *8* | *5.06E-01* | *-4.65* | *3.35E-06* | *6.82E-03* |
| ***Nab2*** | *17937* | *-0.01* | *0.07* | *8.78E-01* | *-0.15* | *0.13* | *8* | *9.83E-01* | *4.58* | *4.72E-06* | *8.23E-03* |
| ***Dph2*** | *67728* | *0.01* | *0.06* | *9.13E-01* | *-0.11* | *0.12* | *8* | *9.87E-01* | *-4.47* | *7.76E-06* | *1.19E-02* |
| ***Wsb1*** | *78889* | *0.15* | *0.12* | *2.00E-01* | *-0.08* | *0.39* | *8* | *7.09E-01* | *4.38* | *1.18E-05* | *1.60E-02* |
| ***Tiparp*** | *99929* | *-0.09* | *0.12* | *4.69E-01* | *-0.33* | *0.15* | *8* | *8.70E-01* | *4.32* | *1.55E-05* | *1.69E-02* |
| ***Fancd2os*** | *70979* | *0.04* | *0.10* | *6.68E-01* | *-0.16* | *0.24* | *7* | *9.35E-01* | *-4.31* | *1.67E-05* | *1.69E-02* |
| ***Unc5a*** | *107448* | *-0.03* | *0.08* | *7.05E-01* | *-0.18* | *0.12* | *8* | *9.45E-01* | *4.30* | *1.72E-05* | *1.69E-02* |
| ***Slc7a2*** | *11988* | *0.01* | *0.06* | *8.59E-01* | *-0.11* | *0.13* | *8* | *9.78E-01* | *-4.28* | *1.84E-05* | *1.69E-02* |
| ***Folr1*** | *14275* | *-0.37* | *0.23* | *1.03E-01* | *-0.81* | *0.08* | *8* | *5.79E-01* | *-4.27* | *1.97E-05* | *1.69E-02* |
| ***Prl4a1*** | *19110* | *-0.35* | *0.25* | *1.71E-01* | *-0.84* | *0.15* | *7* | *6.78E-01* | *-4.23* | *2.32E-05* | *1.69E-02* |
| ***Sympk*** | *68188* | *0.08* | *0.06* | *1.47E-01* | *-0.03* | *0.19* | *7* | *6.48E-01* | *4.23* | *2.32E-05* | *1.69E-02* |
| ***Rhobtb1*** | *69288* | *-0.06* | *0.07* | *4.38E-01* | *-0.20* | *0.09* | *8* | *8.58E-01* | *4.23* | *2.35E-05* | *1.69E-02* |
| ***Zdhhc19*** | *245308* | *-0.39* | *0.31* | *1.98E-01* | *-0.99* | *0.20* | *7* | *7.05E-01* | *-4.16* | *3.13E-05* | *1.99E-02* |
| ***Slc38a1*** | *105727* | *0.05* | *0.03* | *1.13E-01* | *-0.01* | *0.11* | *8* | *5.96E-01* | *4.16* | *3.15E-05* | *1.99E-02* |
| ***Sh2d5*** | *230863* | *-0.03* | *0.06* | *5.88E-01* | *-0.15* | *0.09* | *7* | *9.13E-01* | *4.15* | *3.26E-05* | *1.99E-02* |
| ***Nfil3*** | *18030* | *0.09* | *0.12* | *4.52E-01* | *-0.15* | *0.33* | *8* | *8.62E-01* | *4.12* | *3.80E-05* | *2.12E-02* |
| ***Phlda3*** | *27280* | *0.15* | *0.14* | *2.87E-01* | *-0.13* | *0.43* | *8* | *7.79E-01* | *4.12* | *3.81E-05* | *2.12E-02* |
| ***Ruvbl1*** | *56505* | *0.00* | *0.04* | *9.96E-01* | *-0.08* | *0.08* | *8* | *1.00E+00* | *-4.08* | *4.51E-05* | *2.31E-02* |
| ***Pitpnm2*** | *19679* | *-0.04* | *0.06* | *5.48E-01* | *-0.16* | *0.08* | *8* | *9.04E-01* | *4.08* | *4.54E-05* | *2.31E-02* |
| ***Cx3cr1*** | *13051* | *0.00* | *0.08* | *9.81E-01* | *-0.16* | *0.16* | *8* | *9.98E-01* | *-4.05* | *5.23E-05* | *2.56E-02* |
| ***Srrt*** | *83701* | *0.01* | *0.07* | *9.01E-01* | *-0.13* | *0.15* | *7* | *9.86E-01* | *4.01* | *5.96E-05* | *2.59E-02* |
| ***Csdc2*** | *105859* | *0.13* | *0.10* | *1.68E-01* | *-0.06* | *0.32* | *8* | *6.75E-01* | *4.01* | *6.11E-05* | *2.59E-02* |
| ***Adgrb2*** | *230775* | *0.02* | *0.05* | *6.84E-01* | *-0.08* | *0.13* | *8* | *9.39E-01* | *4.01* | *6.17E-05* | *2.59E-02* |
| ***Gadd45g*** | *23882* | *-0.12* | *0.08* | *1.16E-01* | *-0.28* | *0.03* | *8* | *6.02E-01* | *-4.00* | *6.25E-05* | *2.59E-02* |
| ***Tm4sf1*** | *17112* | *0.00* | *0.09* | *9.81E-01* | *-0.18* | *0.17* | *8* | *9.98E-01* | *4.00* | *6.37E-05* | *2.59E-02* |
| ***Sertad1*** | *55942* | *-0.26* | *0.17* | *1.18E-01* | *-0.59* | *0.07* | *8* | *6.03E-01* | *-3.97* | *7.27E-05* | *2.84E-02* |
| ***Pkhd1l1*** | *192190* | *0.35* | *0.24* | *1.38E-01* | *-0.11* | *0.81* | *8* | *6.34E-01* | *3.95* | *7.66E-05* | *2.84E-02* |
| ***Cdhr1*** | *170677* | *0.29* | *0.26* | *2.59E-01* | *-0.21* | *0.80* | *8* | *7.60E-01* | *3.95* | *7.68E-05* | *2.84E-02* |
| ***Zswim5*** | *74464* | *-0.04* | *0.07* | *5.75E-01* | *-0.17* | *0.09* | *7* | *9.12E-01* | *-3.94* | *7.99E-05* | *2.87E-02* |
| ***Ece1*** | *230857* | *-0.03* | *0.06* | *6.24E-01* | *-0.15* | *0.09* | *7* | *9.21E-01* | *3.91* | *9.25E-05* | *3.23E-02* |
| ***Rgs2*** | *19735* | *-0.06* | *0.05* | *1.78E-01* | *-0.16* | *0.03* | *8* | *6.86E-01* | *-3.90* | *9.55E-05* | *3.24E-02* |
| ***Dock3*** | *208869* | *-0.08* | *0.07* | *2.64E-01* | *-0.22* | *0.06* | *8* | *7.64E-01* | *-3.82* | *1.32E-04* | *4.37E-02* |
| ***Tcn2*** | *21452* | *0.05* | *0.03* | *1.13E-01* | *-0.01* | *0.10* | *8* | *5.96E-01* | *-3.81* | *1.39E-04* | *4.47E-02* |
| ***Mycbpap*** | *104601* | *0.42* | *0.22* | *5.64E-02* | *-0.01* | *0.85* | *8* | *4.80E-01* | *3.79* | *1.50E-04* | *4.71E-02* |

***Table S7. Exploratory meta-analysis results: The full results from the exploratory meta-analysis of antipsychotic effects on the frontal cortex which used data from both rodents and macaques (13,121 genes (10,617 of which had macaque data), 13,107 stable meta-analysis estimates).*** *This .xlsx file includes two worksheets: 1) The worksheet "ColumnDefinitions" provides the definitions for the variables present in each column in "Exploratory_RodentMacaque_MetaAnalysis". 2) The worksheet "Exploratory_RodentMacaque_MetaAnalysis" provides the full meta-analysis results, with each row* *representing the results for one gene, and each column providing either gene annotation or meta-analysis statistical output. The results are ordered by p-value, so that the top rows in the worksheet are the genes with the smallest p-values.*

***Table S8. Exploratory meta-analysis results: All 70 genes consistently upregulated in the frontal cortex following antipsychotic treatment in an exploratory meta-analysis including rodents and macaques (FDR<0.05).*** *Table follows the conventions of* ***Table S3****. Results are ordered by estimated Log2FC (Log2 Fold Change of Antipsychotic treatment vs. Control); a higher Log2FC indicates upregulation of the gene within the frontal cortex following antipsychotic treatment compared to the control group. Given gene symbols are the mouse symbol for the respective gene. Full results for all genes can be found in* ***Table S7****.*

| Gene Symbol | Mouse Entrez Gene ID | Log2FC estimate | SE | pval | CI_lb | CI_ub | # Comparisons | FDR |
|---|---|---|---|---|---|---|---|---|
| Wfikkn1 | 215001 | 0.29 | 0.08 | 4.37E-04 | 0.13 | 0.45 | 10 | 4.90E-02 |
| Gjb1 | 14618 | 0.27 | 0.06 | 4.04E-06 | 0.16 | 0.39 | 8 | 4.41E-03 |
| Nkx6-2 | 14912 | 0.24 | 0.06 | 1.61E-05 | 0.13 | 0.36 | 11 | 8.76E-03 |
| Mfap2 | 17150 | 0.24 | 0.04 | 2.09E-10 | 0.16 | 0.31 | 11 | 2.74E-06 |
| Itgal | 16408 | 0.23 | 0.05 | 2.17E-06 | 0.13 | 0.32 | 10 | 2.84E-03 |
| Igfbp5 | 16011 | 0.22 | 0.05 | 2.27E-05 | 0.12 | 0.32 | 11 | 1.03E-02 |
| Isg15 | 100038882 | 0.21 | 0.06 | 4.07E-04 | 0.10 | 0.33 | 11 | 4.73E-02 |
| Prok1 | 246691 | 0.21 | 0.05 | 7.66E-05 | 0.10 | 0.31 | 7 | 1.90E-02 |
| Calb2 | 12308 | 0.20 | 0.04 | 9.25E-07 | 0.12 | 0.28 | 11 | 1.65E-03 |
| Mroh7 | 381538 | 0.20 | 0.05 | 1.15E-04 | 0.10 | 0.30 | 9 | 2.33E-02 |
| Il13ra1 | 16164 | 0.19 | 0.04 | 5.93E-06 | 0.11 | 0.28 | 7 | 4.86E-03 |
| Gh | 14599 | 0.19 | 0.05 | 1.11E-04 | 0.09 | 0.29 | 8 | 2.30E-02 |
| Slc18a3 | 20508 | 0.18 | 0.05 | 6.85E-05 | 0.09 | 0.27 | 8 | 1.80E-02 |
| Tdo2 | 56720 | 0.18 | 0.04 | 7.78E-06 | 0.10 | 0.26 | 8 | 6.01E-03 |
| Adora2b | 11541 | 0.17 | 0.04 | 4.73E-06 | 0.10 | 0.25 | 11 | 4.58E-03 |
| 1700010I14Rik | 66931 | 0.17 | 0.04 | 3.10E-05 | 0.09 | 0.25 | 8 | 1.16E-02 |
| Osgin1 | 71839 | 0.17 | 0.04 | 2.25E-05 | 0.09 | 0.25 | 11 | 1.03E-02 |
| Gpr84 | 80910 | 0.17 | 0.04 | 4.27E-05 | 0.09 | 0.25 | 8 | 1.27E-02 |
| Slc14a1 | 108052 | 0.17 | 0.04 | 4.88E-06 | 0.10 | 0.24 | 11 | 4.58E-03 |
| Ccdc71 | 72454 | 0.17 | 0.03 | 9.08E-07 | 0.10 | 0.23 | 10 | 1.65E-03 |
| Camk2n2 | 73047 | 0.17 | 0.04 | 1.68E-04 | 0.08 | 0.25 | 11 | 2.92E-02 |
| Wscd1 | 216881 | 0.16 | 0.04 | 2.65E-05 | 0.09 | 0.23 | 11 | 1.12E-02 |

| | | | | | | | | |
|---|---|---|---|---|---|---|---|---|
| ***Slc39a2*** | *214922* | *0.16* | *0.05* | *4.06E-04* | *0.07* | *0.25* | *8* | *4.73E-02* |
| ***Slc1a5*** | *20514* | *0.16* | *0.04* | *1.95E-05* | *0.09* | *0.23* | *8* | *9.83E-03* |
| ***Serinc5*** | *218442* | *0.16* | *0.03* | *2.98E-07* | *0.10* | *0.22* | *11* | *9.77E-04* |
| ***Sfrp5*** | *54612* | *0.15* | *0.04* | *3.83E-04* | *0.07* | *0.24* | *11* | *4.57E-02* |
| ***Chad*** | *12643* | *0.15* | *0.04* | *1.05E-04* | *0.08* | *0.23* | *10* | *2.26E-02* |
| ***F10*** | *14058* | *0.15* | *0.04* | *2.17E-05* | *0.08* | *0.22* | *10* | *1.03E-02* |
| ***Parp12*** | *243771* | *0.15* | *0.04* | *2.80E-04* | *0.07* | *0.23* | *11* | *3.89E-02* |
| ***Spc24*** | *67629* | *0.15* | *0.04* | *9.00E-05* | *0.07* | *0.22* | *10* | *2.11E-02* |
| ***Aspa*** | *11484* | *0.14* | *0.04* | *2.15E-04* | *0.07* | *0.22* | *11* | *3.21E-02* |
| ***Hk2*** | *15277* | *0.14* | *0.04* | *3.21E-04* | *0.07* | *0.22* | *11* | *4.13E-02* |
| ***Megf6*** | *230971* | *0.14* | *0.04* | *1.15E-04* | *0.07* | *0.22* | *11* | *2.33E-02* |
| ***Gpr137b*** | *83924* | *0.14* | *0.04* | *7.33E-05* | *0.07* | *0.21* | *9* | *1.85E-02* |
| ***Gtse1*** | *29870* | *0.14* | *0.04* | *1.08E-04* | *0.07* | *0.21* | *8* | *2.29E-02* |
| ***Mrgprf*** | *211577* | *0.14* | *0.03* | *4.05E-05* | *0.07* | *0.21* | *11* | *1.27E-02* |
| ***Scarf2*** | *224024* | *0.14* | *0.04* | *2.66E-04* | *0.07* | *0.22* | *11* | *3.79E-02* |
| ***Nkain3*** | *269513* | *0.13* | *0.03* | *6.34E-05* | *0.07* | *0.20* | *9* | *1.73E-02* |
| ***Slc29a3*** | *71279* | *0.13* | *0.03* | *4.15E-05* | *0.07* | *0.20* | *11* | *1.27E-02* |
| ***Stx2*** | *13852* | *0.13* | *0.04* | *2.13E-04* | *0.06* | *0.20* | *11* | *3.21E-02* |
| ***Arhgap45*** | *70719* | *0.13* | *0.03* | *1.78E-04* | *0.06* | *0.20* | *11* | *2.92E-02* |
| ***Polr1g*** | *70333* | *0.13* | *0.03* | *1.99E-04* | *0.06* | *0.19* | *7* | *3.11E-02* |
| ***Arhgef10*** | *234094* | *0.12* | *0.03* | *1.02E-04* | *0.06* | *0.19* | *10* | *2.24E-02* |
| ***Rbp7*** | *63954* | *0.12* | *0.03* | *1.82E-04* | *0.06* | *0.19* | *7* | *2.92E-02* |
| ***Nde1*** | *67203* | *0.12* | *0.03* | *9.34E-05* | *0.06* | *0.18* | *11* | *2.13E-02* |
| ***Kit*** | *16590* | *0.12* | *0.03* | *1.00E-05* | *0.06* | *0.17* | *11* | *6.93E-03* |
| ***Cdh3*** | *12560* | *0.12* | *0.03* | *3.56E-04* | *0.05* | *0.18* | *11* | *4.36E-02* |
| ***Fam234a*** | *106581* | *0.12* | *0.03* | *1.56E-04* | *0.06* | *0.18* | *8* | *2.80E-02* |
| ***Cmtm3*** | *68119* | *0.11* | *0.03* | *2.15E-04* | *0.05* | *0.17* | *11* | *3.21E-02* |
| ***Psme2*** | *19188* | *0.11* | *0.02* | *1.71E-06* | *0.07* | *0.16* | *9* | *2.50E-03* |
| ***Cd180*** | *17079* | *0.11* | *0.03* | *4.19E-05* | *0.06* | *0.16* | *11* | *1.27E-02* |
| ***Hnrnpl*** | *15388* | *0.11* | *0.03* | *1.69E-04* | *0.05* | *0.17* | *8* | *2.92E-02* |
| ***Fgfr1*** | *14182* | *0.11* | *0.03* | *4.08E-05* | *0.05* | *0.16* | *11* | *1.27E-02* |
| ***Nrros*** | *224109* | *0.10* | *0.03* | *9.60E-05* | *0.05* | *0.15* | *11* | *2.14E-02* |

| | | | | | | | | |
|---|---|---|---|---|---|---|---|---|
| ***Klhl13*** | *67455* | *0.10* | *0.03* | *3.87E-04* | *0.04* | *0.16* | *10* | *4.57E-02* |
| ***Atad2*** | *70472* | *0.10* | *0.03* | *1.77E-04* | *0.05* | *0.15* | *11* | *2.92E-02* |
| ***Tmem134*** | *66990* | *0.10* | *0.03* | *3.06E-04* | *0.04* | *0.15* | *11* | *4.06E-02* |
| ***Lsm4*** | *50783* | *0.09* | *0.02* | *1.51E-04* | *0.05* | *0.14* | *11* | *2.79E-02* |
| ***Nme3*** | *79059* | *0.09* | *0.03* | *3.65E-04* | *0.04* | *0.14* | *11* | *4.44E-02* |
| ***Rin2*** | *74030* | *0.09* | *0.02* | *2.39E-04* | *0.04* | *0.14* | *11* | *3.52E-02* |
| ***Ctdp1*** | *67655* | *0.09* | *0.02* | *9.40E-05* | *0.04* | *0.13* | *11* | *2.13E-02* |
| ***Slc22a5*** | *20520* | *0.09* | *0.02* | *3.16E-04* | *0.04* | *0.13* | *11* | *4.13E-02* |
| ***Mgat3*** | *17309* | *0.08* | *0.02* | *1.25E-05* | *0.05* | *0.12* | *11* | *7.45E-03* |
| ***Hnrnpa1*** | *15382* | *0.08* | *0.02* | *3.73E-04* | *0.04* | *0.13* | *10* | *4.49E-02* |
| ***Maf1*** | *68877* | *0.08* | *0.02* | *8.66E-05* | *0.04* | *0.12* | *11* | *2.08E-02* |
| ***Rpl27a*** | *26451* | *0.08* | *0.02* | *6.98E-05* | *0.04* | *0.11* | *8* | *1.80E-02* |
| ***Acadvl*** | *11370* | *0.07* | *0.02* | *1.38E-04* | *0.04* | *0.11* | *11* | *2.62E-02* |
| ***Lypla2*** | *26394* | *0.07* | *0.02* | *6.70E-05* | *0.04* | *0.11* | *11* | *1.79E-02* |
| ***Ciao1*** | *26371* | *0.07* | *0.02* | *2.10E-04* | *0.03* | *0.11* | *11* | *3.21E-02* |
| ***Dnajc11*** | *230935* | *0.06* | *0.02* | *2.97E-04* | *0.03* | *0.09* | *11* | *4.03E-02* |

***Table S9. Planned meta-analysis results: All 47 genes consistently downregulated in the frontal cortex following antipsychotic treatment in an exploratory meta-analysis including rodents and macaques (FDR<0.05).*** *Table follows the conventions of **Table S3**. Results are ordered by estimated Log2FC (Log2 Fold Change of Antipsychotic treatment vs. Control); a lower Log2FC indicates downregulation of the gene within the frontal cortex following antipsychotic treatment compared to the control group. Full results for all genes can be found in **Table S7**.*

| ***Gene Symbol*** | ***Mouse Entrez Gene ID*** | ***Log2FC estimate*** | ***SE*** | ***pval*** | ***CI_lb*** | ***CI_ub*** | ***# Comparisons*** | ***FDR*** |
|---|---|---|---|---|---|---|---|---|
| ***Dmkn*** | *73712* | *-0.80* | *0.15* | *1.98E-07* | *-1.10* | *-0.50* | *7* | *8.66E-04* |
| ***Ms4a8a*** | *64381* | *-0.25* | *0.04* | *6.28E-09* | *-0.33* | *-0.16* | *7* | *4.12E-05* |
| ***Dglucy*** | *217830* | *-0.21* | *0.06* | *3.39E-04* | *-0.33* | *-0.10* | *7* | *4.23E-02* |
| ***Nr0b1*** | *11614* | *-0.17* | *0.05* | *4.23E-04* | *-0.27* | *-0.08* | *7* | *4.78E-02* |
| ***Fabp4*** | *11770* | *-0.17* | *0.04* | *5.36E-05* | *-0.25* | *-0.09* | *7* | *1.50E-02* |
| ***Cth*** | *107869* | *-0.16* | *0.04* | *1.67E-04* | *-0.24* | *-0.07* | *11* | *2.92E-02* |

| | | | | | | | | |
|---|---|---|---|---|---|---|---|---|
| ***Or51e2*** | *170639* | *-0.15* | *0.04* | *3.24E-04* | *-0.24* | *-0.07* | *8* | *4.13E-02* |
| ***Pus7l*** | *78895* | *-0.15* | *0.04* | *1.67E-05* | *-0.22* | *-0.08* | *10* | *8.76E-03* |
| ***Proser3*** | *333193* | *-0.15* | *0.04* | *2.64E-04* | *-0.23* | *-0.07* | *9* | *3.79E-02* |
| ***Cdhr5*** | *72040* | *-0.15* | *0.04* | *4.21E-04* | *-0.23* | *-0.07* | *7* | *4.78E-02* |
| ***Opn3*** | *13603* | *-0.15* | *0.04* | *3.76E-05* | *-0.22* | *-0.08* | *11* | *1.27E-02* |
| ***Trib1*** | *211770* | *-0.14* | *0.04* | *1.82E-04* | *-0.22* | *-0.07* | *11* | *2.92E-02* |
| ***Kcnh5*** | *238271* | *-0.14* | *0.04* | *1.22E-04* | *-0.21* | *-0.07* | *11* | *2.42E-02* |
| ***Tatdn1*** | *69694* | *-0.14* | *0.03* | *5.34E-06* | *-0.19* | *-0.08* | *10* | *4.67E-03* |
| ***Ar*** | *11835* | *-0.13* | *0.03* | *1.24E-04* | *-0.19* | *-0.06* | *11* | *2.43E-02* |
| ***Tmem254*** | *66039* | *-0.12* | *0.03* | *1.45E-04* | *-0.19* | *-0.06* | *7* | *2.71E-02* |
| ***Susd2*** | *71733* | *-0.12* | *0.03* | *2.77E-04* | *-0.18* | *-0.06* | *8* | *3.89E-02* |
| ***Ncr1*** | *17086* | *-0.12* | *0.03* | *3.44E-05* | *-0.18* | *-0.06* | *7* | *1.25E-02* |
| ***Slc35g2*** | *245020* | *-0.11* | *0.02* | *3.46E-06* | *-0.16* | *-0.06* | *9* | *4.13E-03* |
| ***Snx16*** | *74718* | *-0.11* | *0.03* | *1.73E-04* | *-0.16* | *-0.05* | *11* | *2.92E-02* |
| ***Chic1*** | *12212* | *-0.11* | *0.02* | *1.01E-06* | *-0.15* | *-0.06* | *10* | *1.65E-03* |
| ***Nudt7*** | *67528* | *-0.11* | *0.03* | *4.17E-04* | *-0.16* | *-0.05* | *11* | *4.78E-02* |
| ***Ttc39b*** | *69863* | *-0.10* | *0.02* | *4.51E-05* | *-0.15* | *-0.05* | *11* | *1.32E-02* |
| ***Wdpcp*** | *216560* | *-0.10* | *0.02* | *2.88E-05* | *-0.14* | *-0.05* | *11* | *1.14E-02* |
| ***Mrps30*** | *59054* | *-0.10* | *0.02* | *5.91E-07* | *-0.14* | *-0.06* | *11* | *1.55E-03* |
| ***Lrrc8c*** | *100604* | *-0.10* | *0.02* | *2.97E-05* | *-0.14* | *-0.05* | *11* | *1.15E-02* |
| ***Polr3c*** | *74414* | *-0.10* | *0.03* | *3.36E-04* | *-0.15* | *-0.04* | *11* | *4.23E-02* |
| ***Errfi1*** | *74155* | *-0.09* | *0.02* | *1.35E-05* | *-0.14* | *-0.05* | *11* | *7.68E-03* |
| ***Sdhaf3*** | *71238* | *-0.09* | *0.03* | *2.82E-04* | *-0.14* | *-0.04* | *11* | *3.89E-02* |
| ***Fam13c*** | *71721* | *-0.09* | *0.02* | *9.57E-06* | *-0.13* | *-0.05* | *10* | *6.93E-03* |
| ***Syt4*** | *20983* | *-0.09* | *0.02* | *5.14E-05* | *-0.14* | *-0.05* | *11* | *1.47E-02* |
| ***Cwc22*** | *80744* | *-0.09* | *0.02* | *2.75E-05* | *-0.13* | *-0.05* | *10* | *1.13E-02* |
| ***Osbpl8*** | *237542* | *-0.09* | *0.02* | *2.39E-05* | *-0.13* | *-0.05* | *11* | *1.04E-02* |
| ***Zdhhc5*** | *228136* | *-0.09* | *0.02* | *8.70E-05* | *-0.13* | *-0.04* | *11* | *2.08E-02* |
| ***Psmd12*** | *66997* | *-0.09* | *0.02* | *1.19E-05* | *-0.13* | *-0.05* | *11* | *7.45E-03* |
| ***Tasp1*** | *75812* | *-0.08* | *0.02* | *1.79E-04* | *-0.13* | *-0.04* | *11* | *2.92E-02* |

| | | | | | | | | |
|---|---|---|---|---|---|---|---|---|
| ***Gtf3c3*** | *98488* | *-0.08* | *0.02* | *2.45E-04* | *-0.13* | *-0.04* | *10* | *3.57E-02* |
| ***Phf14*** | *75725* | *-0.08* | *0.02* | *2.98E-04* | *-0.13* | *-0.04* | *11* | *4.03E-02* |
| ***Prickle1*** | *106042* | *-0.08* | *0.02* | *3.18E-04* | *-0.12* | *-0.04* | *11* | *4.13E-02* |
| ***Pwp2*** | *110816* | *-0.08* | *0.02* | *1.56E-04* | *-0.12* | *-0.04* | *11* | *2.80E-02* |
| ***Rbm12*** | *75710* | *-0.08* | *0.02* | *1.31E-04* | *-0.12* | *-0.04* | *7* | *2.52E-02* |
| ***Hace1*** | *209462* | *-0.08* | *0.02* | *3.01E-04* | *-0.12* | *-0.03* | *11* | *4.03E-02* |
| ***Tfb2m*** | *15278* | *-0.07* | *0.02* | *1.96E-04* | *-0.11* | *-0.03* | *11* | *3.10E-02* |
| ***Dctn4*** | *67665* | *-0.07* | *0.02* | *4.20E-05* | *-0.11* | *-0.04* | *10* | *1.27E-02* |
| ***Aptx*** | *66408* | *-0.07* | *0.02* | *4.00E-05* | *-0.11* | *-0.04* | *11* | *1.27E-02* |
| ***Camsap2*** | *67886* | *-0.07* | *0.02* | *1.23E-05* | *-0.10* | *-0.04* | *11* | *7.45E-03* |
| ***Ipo7*** | *233726* | *-0.06* | *0.02* | *3.48E-04* | *-0.09* | *-0.03* | *11* | *4.30E-02* |

***Table S10: Planned Rodent Meta-Analysis: Non-directional fast Gene Set Enrichment Analysis (fGSEA) results (10,147 gene sets).*** *This .xlsx file includes two worksheets: 1) The worksheet "ColumnDefinitions" provides the definitions for the variables present in each column in "RodentMetaAnalysis_nondirectional_GSEA". 2) The worksheet "RodentMetaAnalysis_nondirectional_GSEA" provides the full fGSEA results, with each row representing the results for one gene set, and each column providing the fGSEA statistical output. The results are ordered by p-value, so that the top rows in the worksheet are the gene sets with the smallest p-values.*

***Table S11: Planned Rodent Meta-Analysis: Directional fast Gene Set Enrichment Analysis (fGSEA) results (10,147 gene sets).*** *This .xlsx file includes two worksheets: 1) The worksheet "ColumnDefinitions" provides the definitions for the variables present in each column in "RodentMetaAnalysis_DirectionalGSEA". 2) The worksheet "RodentMetaAnalysis_DirectionalGSEA" provides the full fGSEA results, with each row representing the results for one gene set, and each column providing the fGSEA statistical output. The results are ordered by p-value, so that the top rows in the worksheet are the gene sets with the smallest p-values.*

***Table S12: Exploratory Rodent and Macaque Meta-Analysis: Non-directional fast Gene Set Enrichment Analysis (fGSEA) results (10,417 gene sets).*** *This .xlsx file includes two worksheets: 1) The worksheet "Column Definitions" provides the definitions for the variables present in each column in "Exploratory_RodentMacaque_GSEA_NonDirectional". 2) The worksheet "Exploratory_RodentMacaque_GSEA_NonDirectional" provides the full fGSEA results, with each row representing the results for one gene set, and each column providing the fGSEA statistical output. The results are ordered by p-value, so that the top rows in the worksheet are the gene sets with the smallest p-values.*

***Table S13: Exploratory Rodent and Macaque Meta-Analysis: Directional fast Gene Set Enrichment Analysis (fGSEA) results (10,417 gene sets).*** *This .xlsx file includes two worksheets: 1) The worksheet "Column Definitions" provides the definitions for the variables present in each column in "Exploratory_RodentMacaque_GSEA_Directional". 2) The worksheet "Exploratory_RodentMacaque_GSEA_Directional" provides the full fGSEA results, with each row representing the results for one gene set, and each column providing the fGSEA statistical output. The results are ordered by p-value, so that the top rows in the worksheet are the gene sets with the smallest p-values.*

### Supplemental Figures

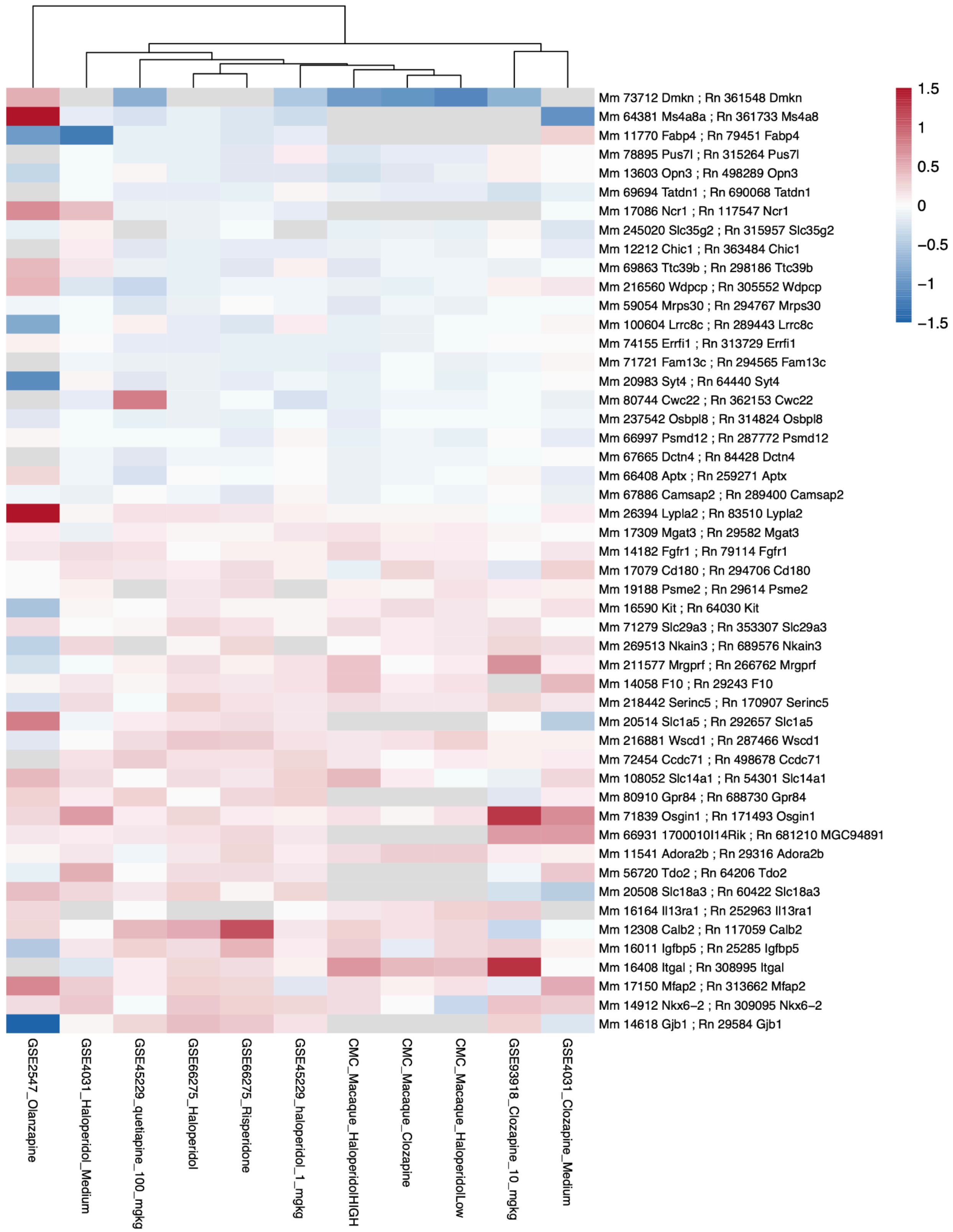

***Figure S1. A hierarchically-clustered heatmap illustrating the antipsychotic vs. control effect sizes across all group comparisons for each of the top 50 differentially expressed genes (DEGs) in the exploratory meta-analysis using both rodent and macaque data.*** *The heatmap displays the antipsychotic vs. control effect sizes (Log(2) fold changes or Log2FCs) for each of the 50 differentially expressed genes identified through the meta-analysis as significantly differentially expressed (FDR<0.05). Results are scaled and color-coded by log2FC, ranging from -1.5 to +1.5 (blue to red; downregulation to upregulation). Rows represent individual genes, annotated with the ortholog mouse (Mm) and rat (Rn) Entrez ids and gene symbols. Each column corresponds to a specific treatment vs. control comparison, identified by Gene Expression Omnibus dataset id (GSE…) and treatment used. Hierarchical clustering is applied and used to group columns with similar expression profiles.*
